# Phase Separation and Aggregation of α-Synuclein Diverge at Different Salt Conditions

**DOI:** 10.1101/2024.03.01.582895

**Authors:** Rebecca Sternke-Hoffmann, Xun Sun, Andreas Menzel, Miriam Dos Santos Pinto, Urtė Venclovaitė, Michael Wördehoff, Wolfgang Hoyer, Wenwei Zheng, Jinghui Luo

## Abstract

The coacervation and structural rearrangement of the protein alpha-synuclein (αSyn) into cytotoxic oligomers and amyloid fibrils are considered pathological hallmarks of Parkinson’s disease. While aggregation is recognized as the key element of amyloid diseases, liquid-liquid phase separation (LLPS) and its interplay with aggregation have gained increasing interest. Previous work showed that factors promoting or inhibiting amyloid formation have similar effects on phase separation. Here, we provide a detailed scanning of a wide range of parameters including protein, salt and crowding concentrations at multiple pH values, revealing different salt dependencies of aggregation and phase separation. The influence of salt on aggregation under crowded conditions follows a non-monotonic pattern, showing increased effects at medium salt concentrations. This behavior can be elucidated through a combination of electrostatic screening and salting-out effects on the intramolecular interactions between the N-terminal and C-terminal regions of αSyn. By contrast, we find a monotonic salt dependence of phase separation due to the intermolecular interaction. Furthermore, we observe the time evolution of the two distinct assembly states, with macroscopic fibrillar-like bundles initially forming at medium salt concentration but subsequently converting into droplets after prolonged incubation. The droplet state is therefore capable of inhibiting aggregation or even dissolving the aggregates through a variety of heterotypic interactions, thus preventing αSyn from its dynamically arrested state.

## Introduction

Parkinson’s disease (PD) is a prevalent neurodegenerative disorder that has seen a rise in incidence over the last three decades. It is characterized by the loss of dopaminergic neurons in the substanita nigra, resulting in motor and non-motor symptoms(1). As life expectancy continues to increase globally, PD has become a significant challenge to global health(2). The pathological hallmark of PD is the coacervation and structural rearrangement of the protein α-synuclein (αSyn) into cytotoxic oligomers, amyloid fibrils, and ultimately into Lewy bodies(3, 4). These misfolded, aggregated, and over-accumulated proteins are not able to be cleared by the protein quality control machinery, leading to impaired cellular functions, cell death, and neurodegeneration. Despite its pathological significance in PD, αSyn is also known to be involved in several physiological functions, including neurotransmitter release, axonal transport, and synaptic vesicle trafficking(5). However, the precise role of αSyn in these processes remains unclear.

αSyn is an intrinsically disordered protein (IDP) consisting of three distinct domains with 140 amino acids: the highly conserved amphipathic N-terminal domain (NTD), the hydrophobic non-amyloid component (NAC) region, and the flexible C-terminal tail (CTT)(6). The hydrophobic side chains of the NAC region of αSyn are proposed to be primary drivers of αSyn aggregation building the fibril core(7). Part of the NTD and most of the CTT extend from the fibril core like a polymer brush(8, 9). However, most familial missense mutations (e.g. A53T, E46K, and A30P) are located in the α-helical NTD(10–12) and a short motif inside NTD has been recognized to be critical for its aggregation(13). Experimental evidence demonstrated long-range interactions between CTT and the NTD or NAC(6, 14), which might protect the NAC region and inhibits aggregation (15–17).

Due to the disordered nature and conformational flexibility of αSyn, cellular and environmental factors can play a major role in its relevant disease pathogenesis and aggregation. With lower levels of sodium and chloride in serum, Parkinson’s patients are more likely to develop dyskinesia(18). In contrast, salt concentration increase *in vitro* can effectively reduce attractive interactions between the net positively charged NTD and negatively charged CTT, leading to increased solvent exposure and aggregation of the NAC region. Increasing salt concentration promotes αSyn aggregation at pH 7(19). Reducing pH below 6 significantly increases the rate of secondary nucleation but only slightly enhances the rate of elongation(20). However, secondary nucleation is retarded at pH 5.5 as salt concentration increases(17), suggesting that the interplay between salt and pH require specific consideration of the aggregation pathway. Moreover, macromolecular crowding, mimicking intracellular conditions (21) can accelerate αSyn aggregation primarily through volume exclusion, despite the reduction in diffusion resulting from increased viscosity (22). Crowding reagents (crowders) with longer chain length promote faster aggregation than those with shorter chain length(23). The influence of macromolecular crowding reagents is evident across primary nucleation, fiber elongation, and secondary nucleation processes (24, 25).

Recent studies demonstrated the ability of αSyn to undergo liquid-liquid phase separation (LLPS) both *in vitro* and *in vivo*(26, 27). A variety of disordered proteins have been observed to form these liquid-droplet like biomolecular condensates, which compartmentalize biomolecules (e.g. proteins and nucleic acids) into intracellular membraneless organelles, covering a large number of compositions, structural features and functions(28–33). Besides αSyn, many other amyloid-forming proteins such as Tau(34), FUS(35, 36), TDP-43(37), polyQ(38), IAPP(39), hnRNPA1(40), and hnRNPA2(41) are also known to undergo LLPS through their intrinsically disordered and/or low-complexity domains. The presence of biological condensates can drastically affect the aggregation process. In many of these examples, the liquid droplets can effectively increase the local concentration of the protein of interest and possibly promote conformational changes within the disordered region of the proteins, leading to a gel-like and subsequent solid-like state(42). Condensation through LLPS may increase the nucleation rate into amyloid fibrils and might be the triggering factor ultimately to the development of disease(34). It would then be interesting to investigate the relation between phase separation and aggregation of αSyn, which can possibly shed light upon other LLPS-enabled and fibril-forming proteins with similar properties.

It has been also suggested that the liquid droplet state acts as the precursor of αSyn nucleation (26). Upon varying the cellular and environmental factors, the tendencies of phase separation and aggregation often change similarly. For instance, increasing the molecular crowding and decreasing the pH to 5.4 lead to a decreased critical protein concentration for LLPS(26). Both changes accelerate αSyn aggregation(20, 22). Similarly, increasing salt concentration promotes LLPS(43) as also observed in aggregation(19). Furthermore, PD-associated factors, like Cu^2+^, Fe^3+^, and liposomes, accelerate droplet formation and the molecules inside the droplets show a higher rigidity(26). Both calcium and manganese ions promote droplet formation as well as amyloid aggregation(44, 45). The CTT truncated αSyn has also been found to accelerate WT αSyn phase separation and aggregation(46). One possible explanation for several of these factors lies in the ability to induce an elongated conformation, featuring a greater solvent-exposed NAC surface, which in turn facilitates an accelerated progression of both LLPS and aggregation. Lowering pH, elevating salt concentrations, or truncating the CTT presumably diminish the intramolecular interactions between the CTT and the NTD and NAC regions. This shift could effectively transform the collapsed conformation of αSyn towards more elongated conformations, facilitating the requisite protein-protein interactions crucial for LLPS and aggregation. Indeed, elongated and more hairpin-like conformation has been observed upon LLPS providing one such evidence(47). Yet, when contemplating functional significance, phase separation is commonly linked to intracellular functions, whereas aggregation, particularly in the case of αSyn, is associated with pathological diseases. It becomes challenging to conceive that the sequence determinants of phase separation and aggregation have co-evolved. In fact, a recent study proposed that αSyn condensation may both expedite and hinder its aggregation, contingent on the localization of aggregates within the droplets. This implies that LLPS may not necessarily be correlated with aggregation in all circumstances (48). Such cases in which phase separation does not necessarily develop at the same condition as aggregation have started to be observed recently in a few other examples, such as FUS(49), TDP43(50), and Aβ42(51).

In the present study, we focus on investigating the initiation and evolution of phase separation and aggregation of αSyn under varied salt, pH, and crowding conditions which are prepared by using the protein crystallization robotic dispenser. The correlation between phase separation and aggregation of αSyn is observed by protein crystallization robotic imaging and fibrillation kinetics. To provide a crowded environment, aggregation and phase separation of αSyn was studied in the presence of PEG400, a medium sized crowder which is known to affect the conformational preference and intrachain dynamics of intrinsically disordered proteins (52). Through systematically manipulating various factors for αSyn assembly in simulations and experiments, we disentangle molecular and sequence determinants of phase separation and aggregation at mesoscales. Our work can help deepen our understanding of the underlying molecular pathogenesis of Parkinson’s disease and potentially contribute to the development of new therapies.

## Results

### Regulation of αSyn aggregation by intramolecular electrostatic interactions between NTD and CTT

We first take a closer look at the sequence composition of αSyn in Fig. 1A. αSyn is with ∼27.9% charged amino acids, slightly higher than 27.7% for IDPs from DisProt database(53). Such a charge content suggests the charged amino acids could play a significant role in determining the conformational preference of the protein. A more detailed inspection suggests αSyn has a net charge of –9 at physiological pH and these charges are not uniformly distributed along the sequence. The three distinct domains of αSyn are the N-terminal domain (NTD, residue 1 to 60), the hydrophobic non-amyloid-β component (NAC, residue 61 to 95) region, and the flexible C-terminal tail (CTT, residue 96 to 140). The NTD is slightly positively charged with a net charge of +4 and the CTT is highly negatively charged with a net charge of -12. There is limited charged amino acids in the NAC, suggesting limited electrostatic interactions between NAC and the other two parts of the chain. At physiological pH, we expect long-range intramolecular interactions between the NTD and CTD, making NAC more solvent buried and less prone to aggregate (Fig. 1B). When reducing pH, the net charge of CTT starts to decrease, leading to a more elongated conformation and more solvent exposed NAC for aggregation (Fig. 1C). The C-terminal truncated protein (ΔCTT variant with residues from 1 to 108, Fig. 1D) should behave similarly as the full-length αSyn at low pH without any NTD and CTT interactions.

**Figure 1.**
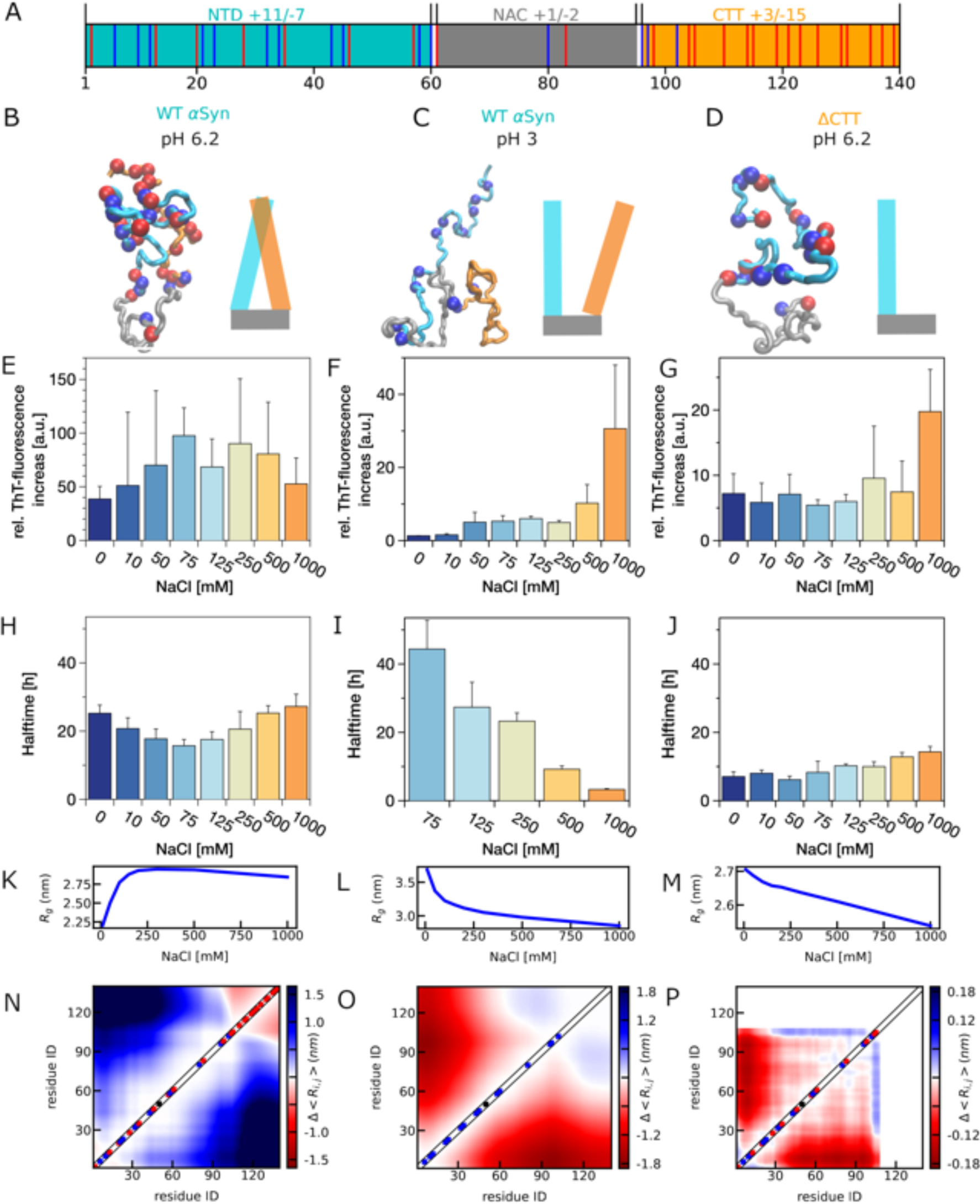
A) Distribution of charged amino acids in αSyn sequence, with positively charged amino acids highlighted in blue and negatively charged ones highlighted in red. The N-terminal domain (NTD, residue 1 to 60) is colored in cyan, non-amyloid-β component (NAC, residue 61 to 95) colored in grey and the C-terminal tail (CTT, residues 96-140) colored in orange. B, C, D) Representative conformations of αSyn variants including full length αSyn at pH 6.2 (B), at pH 3.0 (C), and ΔCTT variant at pH 6.2 (D). E, F, G) Salt-dependent aggregation of these αSyn variants assessed by ThT fluorescence intensity increase. 15 µM αSyn was incubated in the presence of 20% PEG400 at 37°C under shaking conditions with increasing salt concentrations. H, I, J) The corresponding half times for the aggregation of these αSyn variants. K, L, M) Radius of gyration (*R_g_*) of these αSyn variants from the coarse-grained molecular dynamics simulations. N, O, P) The difference between the pairwise distance maps at 100 mM and 10 mM salt concentrations from the simulations.

To confirm the role of CTT and more specifically the electrostatic interactions on αSyn aggregation, we performed ThT-incorporation experiments for two αSyn variants under varied salt concentrations and pH conditions. We present their aggregation in Fig. 1E to G in the presence of molecular crowders (i.e. 20% PEG400) and further investigate other crowding conditions in the next section. It was observed previously that at neutral pH under quiescent conditions homogeneous primary nucleation and secondary processes are very slow(20). To enhance primary nucleation and facilitate fragmentation, we performed the aggregation kinetics experiments under shaking conditions at pH 6.2 (raw data in Fig. S1). The presence of amyloid fibrils could also be verified under shaking and quiescent conditions using atomic force microscopy (AFM) (Fig. S2). While shaking led to a more homogeneous length distribution of the fibrils, we observed a mixture of long and short fibrils after quiescent incubation and furthermore the presence of small oligomers particularly at 1000 mM salt. AFM revealed circular structures ranging from a few nanometers to 1-2 micrometers in size, associated with amyloid fibrils within the 75 µM αSyn sample incubated under quiescent conditions with 500 mM salt. These might be liquid droplets from the liquid-liquid phase separation of αSyn, which were further investigated in the next section.

First, we look at the salt-dependent aggregation close to neutral pH (pH 6.2). We observe salt-induced aggregation at low salt concentration (<250 mM) as was reported by others(19). This can be explained by the reducing intramolecular electrostatic interactions between the NTD and CTD, resulting in more solvent exposed NAC for aggregation (Fig. 1E and H). However, we find the aggregation kinetics are slowed down when increasing the salt concentration above 250 mM. Such a nonmonotonic salt dependence cannot be interpreted solely by electrostatic screening. Since ThT affinity to amyloid fibrils can be salt dependent (54), we have also measured the free monomer concentration at the supernatant and observed the same non-monotonic salt dependence (Fig. S3).

To find a molecular level understanding, we utilized a recently developed coarse-grained simulation model for investigating the conformational preference of αSyn. In order to capture the behavior of αSyn at very high salt concentrations, we used an extension of the original HPS model(55), the HPS-salt model(56), which takes into account the salting-out effect. The salting-out effect was shown to be necessary to capture the salt-dependent conformational change of IDPs, especially for ionic strength above 500 mM. There is one free parameter, the interaction strength ε of the pairwise interaction term, which can be adjusted according to the experimental measurement of the size of a specific IDP. We observed ε=0.13 kcal/mol as the best to capture the existing Förster resonance energy transfer (FRET) measurement of αSyn without crowding (57) as shown in Fig. S4. The same parameter was found for Aβ40 in a previous work (58). To simulate αSyn at 20% and 25% of PEG400 conditions, we increased the interaction strength between pair of amino acids from 0.13 kcal/mol to 0.15 and 0.17 kcal/mol, respectively.

As shown in Fig. 1K, the radius of gyration (*R_g_*) of αSyn from a single-chain simulation (ε=0.15 kcal/mol) first increases and then decreases when increasing the salt concentrations. The turning point of ∼300 mM is in close agreement with the experimental observed turning point of 250 mM for the salt-dependent aggregation behavior, suggesting an expanded conformation and probably more solvent exposed NAC is a good indicator of its aggregation, consistent with the previous literature(47). To further confirm our observation from the simulation, we conducted small-angle X-ray scattering (SAXS) experiments on αSyn under varying salt concentrations, in the presence or absence of 20% PEG400. The presence of possible aggregates, oligomers, and monomers in the solution posed a challenge in confidently estimating the size of the monomers. In an attempt to address this complexity, as illustrated in Fig. S5 and S6, we strategically separated the Guinier regions of the dataset, leveraging the size differences between aggregates and monomers. This allowed us to obtain an estimate of the radius of gyration (*R_g_*) as a function of salt concentrations. Despite observing larger *R_g_* values compared to CG simulations for all cases, possibly attributed to high protein concentrations in experiments and the influence of assembly states other than monomers, we consistently observed a qualitatively consistent nonmonotonic salt dependence for αSyn, both with and without PEG400.

We further look at the interactions that contribute to the salt-dependent size of αSyn in Fig. 1N and find that the distance variation between the two ends of the protein, corresponding to the NTD-CTT interactions, can explain size variation. The major reason for the further collapse above 300 mM is due to the salting-out effect driven by the hydrophobic amino acids(56, 59), which is independent to the electrostatic screening driven by the charged amino acids. The interplay between the electrostatic screening and the salting-out effect then provides such a nonmonotonic salt-dependence for the size of αSyn.

On the other hand, reducing pH is expected to change the salt dependence of αSyn at a low salt concentration, since it effectively neutralizes the negatively charged amino acids at CTT. At pH 3 as shown in Fig. F and I, αSyn demonstrated the fastest kinetics and highest fluorescence increase at a high salt concentration. The non-monotonic salt-dependence disappears: the salt dependence at a high salt concentration stays the same due to the same salting-out effect; the salt dependence at low salt concentration at pH 3 is opposite to that at pH 6.2, since salt screens the net repulsive interactions of NTD instead of the attractive interactions between NTD and CTT. Shown in Fig. 1L, the observation of a monotonically reducing size of αSyn when increasing salt concentrations in simulations is consistent with our experimental data. Due to the net repulsive intramolecular interactions between the positively charged NTD and now slightly positively charged CTT at pH 3 in contrast to the negatively charged CTT at pH 6.2, increasing salt concentration and electrostatic screening result in weaker repulsive interactions and smaller distances between NTD and CTT (Fig. 1O). Again, the salt-dependent aggregation behavior of αSyn fully correlates with and can be explained by the single-chain size of and intramolecular interactions of αSyn in simulations.

A variant of αSyn, ΔCTT, has been previously introduced to understand the role of CTT in its aggregation (14). Here we expect that without CTT interaction with NTD and NAC, aggregation will become much faster. In addition, the non-monotonic salt-dependence as seen in WT αSyn at pH 6.2 should disappear similar to the pH 3 case. As shown in Fig.1J, the ΔCTT (variant with residues from 1 to 108) aggregates faster than the full-length αSyn as expected. However, both ThT intensity and kinetics (Fig. 1J and G) display a much weaker monotonic salt dependence in comparison to the WT αSyn at pH 3. We also confirm from the simulation there is a much weaker salt-dependent size variation in contrast to that of WT αSyn at pH 3 (Fig. 1M). The *R_g_* reduced by about 30% for WT αSyn at pH 3 when increasing the salt concentration from 10 mM to 2000 mM, but only about 12% for ΔCTT at pH 6.2. A closer look at the distance variation (Fig. 1M and P) suggests that without CTT, the size of the protein with only NTD and NAC will not change dramatically when varying the salt. Therefore, the solvent exposure of NAC and the corresponding aggregation of αSyn ΔCTT is only weakly affected by the salt.

We have performed the salt and pH-dependent aggregation measurements without the crowding agents as shown in Fig. S7. The crowding accelerates the aggregation in all cases as observed previously(24, 25). The salt dependent behaviors are qualitatively similar for WT αSyn at pH 3 and ΔCTT at pH 6.2. However, for WT αSyn at pH 6.2, we saw a more monotonic trend for both the ThT fluorescence increase and the aggregation half time, suggesting that the salt-dependent aggregation mechanism could vary in the presence and absence of crowders(24). Specifically at high salt concentrations, the aggregation is not inhibited by increased intramolecular NTD-CTT interactions as observed in a crowded environment, since salt can also enhance intermolecular-NAC interactions through the salting-out effect(59). The interplay between these intra- and intermolecular interactions will further be investigated in the next section.

Through a series of experiments and simulations of two different variants of αSyn under varied salt and pH conditions, a robust correlation has emerged between the observed experimental aggregation behavior and the simulated size of αSyn under crowded conditions. Specifically, in the presence of CTT, as seen in the full-length αSyn, a more expanded αSyn tends to aggregate faster and more readily. The size of αSyn is primarily influenced by intramolecular electrostatic interactions between charged amino acids within NTD and CTT. The key molecular driving force behind αSyn aggregation is the inhibition of interactions between the net positive NTD and the net negative CTT. However, when CTT is not in play, such as in αSyn at low pH or ΔCTT, electrostatic screening by the net positive NTD propels αSyn aggregation.

### Multivariate effect of crowding and salt on αSyn phase separation

Recent studies proved the ability of αSyn to form liquid-like droplets *in vitro* in the presence of molecular crowders(26). Without crowding, we cannot observe αSyn LLPS at the investigated protein concentrations. In order to investigate the multivariate effect of both crowding and salt concentration on αSyn LLPS, we first took three crowding conditions (i.e. 18%, 20%, and 25% PEG400) which are close to the crowding conditions used in αSyn aggregation. Compared to previous publications investigating LLPS formation at pH 7.4, we examine droplet formation at slightly more acidic pH values (pH 6.2 and pH 7). Unlike the aggregation experiments, the LLPS formation was investigated under quiescent conditions and because of technical reasons, the experiments were conducted at 20°C. In order to confirm that the salt-dependence of aggregation was not temperature dependent, we conducted aggregation experiments at 25°C under quiescent conditions at a higher protein concentration and observed a similar trend (Fig. S8). However, at certain conditions the aggregation would be too slow for a reasonable time frame.

A small droplet (600 nl) of αSyn solution was mixed with the buffer solution 1:1 into a well of a MRC 3-well plate to achieve the desired PEG and NaCl concentration. Using the RockImager microscope, we achieved a long investigation time frame for LLPS formation unlike aggregation. As shown in Fig. 2A after 29 days of incubation, we could not observe any macroscopic droplet formation in the presence of 18% PEG400. Droplets are formed in the presence of 20% PEG400 at high salt concentrations (≥250 mM NaCl). At 25% PEG, droplets can be observed at all salt concentrations (≥50 mM NaCl). We did all the experiments at a monomer concentration of 15 µM, in contrast to the previous studies, showing a critical concentration of higher than 50 µM(46) and 200 µM(26) at neutral pH. These observations suggest that both crowding and salt concentration can induce the αSyn LLPS. However, both the trends are monotonic, different from the non-monotonic salt-dependence seen in αSyn aggregation.

**Figure 2.**
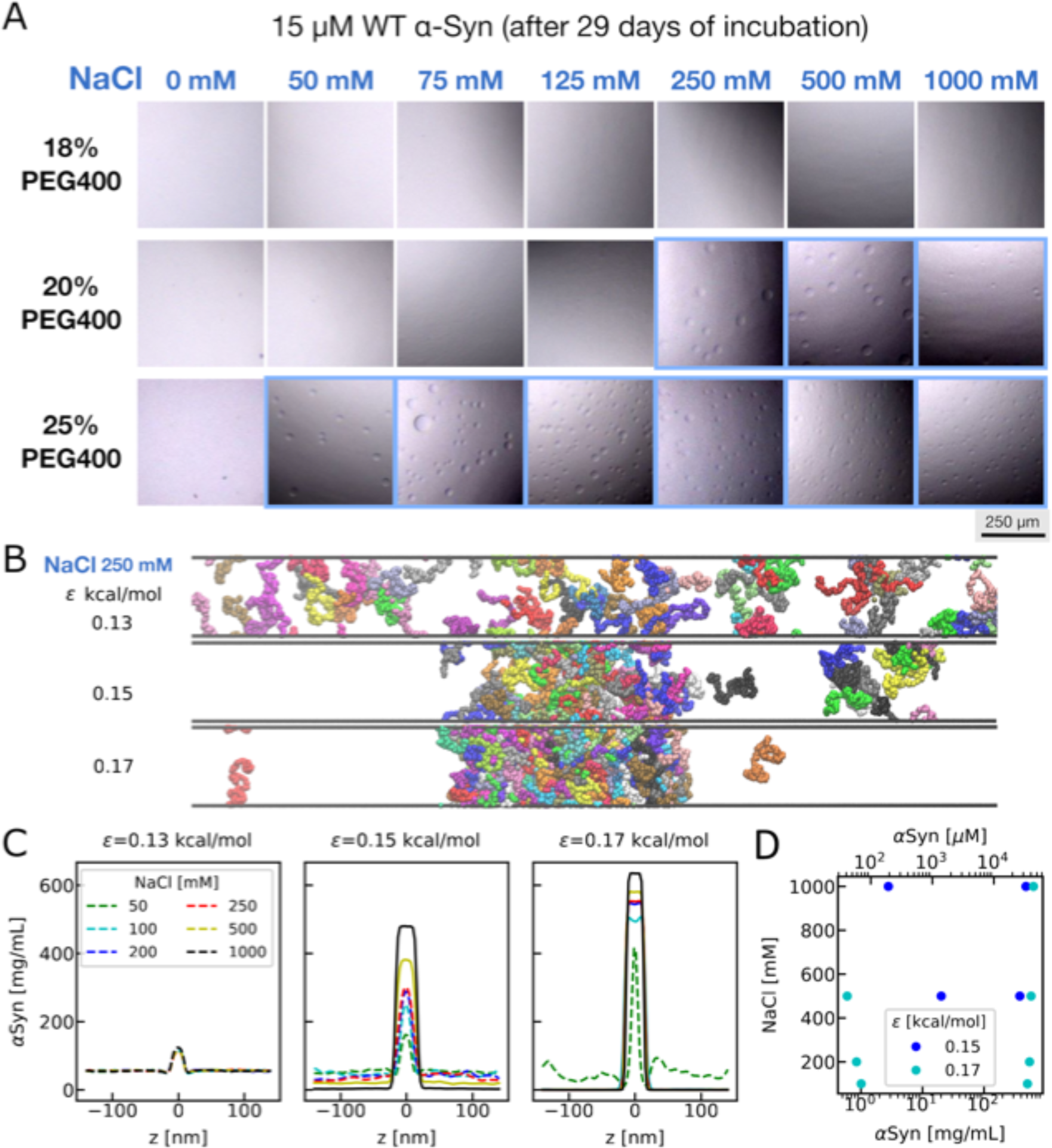
(A) Micrographs (500 x 500 µm) of liquid droplet formation of 15 µM WT αSyn at pH 6.2 at different salt concentrations (0, 50, 75, 125, 500 and 1000 mM NaCl) in the presence of 18%, 20%, and 25% PEG400. (B) Representative configurations from the coarse-grained slab simulations at 250 mM NaCl with three different interaction strengths. (C) Density profiles from slab simulations at different interaction strengths and salt conditions. The density profile in which a slab is well defined for determining the coexistence densities is shown in the solid lines and the one in which the slab is broken is shown in the dashed lines. (D) Phase diagrams obtained using the density profiles in (C).

We aim to gain a molecular-level comprehension of how crowders and salts influence αSyn liquid-liquid phase separation (LLPS) through our coarse-grained simulation model, which has been employed previously to elucidate αSyn aggregation in this study. Based on the polymer theory(60) a homopolymer’s single-chain behavior (e.g. theta solvent temperature at which the protein behaves like a random coil(61), or size of the protein(62)) positively correlates with the LLPS behavior (e.g. critical temperature). However, in this case, we observe a non-monotonic salt dependence for the size of αSyn, contrasting with the monotonic salt dependence for LLPS. This discrepancy suggests a breakdown in the correlation between the single-chain behavior and LLPS in the context of αSyn. This could be understood by the specific NTD and CTT interactions within αSyn, which deviates from a homopolymer(63). We, then, utilized a previously tested slab simulation method(55, 64, 65) to directly simulate the coexistence of the low- and high-density phases of αSyn. We prepared an initial setup in which 100 chains of αSyn were clustered into a slab conformation at the center of the box. We then performed the simulation at multiple conditions with a special shape of the simulation box in which the z-axis is much longer than the x- and y-axis.

To consider the crowding conditions implicitly in the coarse-grained simulations, we increase the amino acid interaction strength. As shown in Fig. 2B, we show representative conformations from simulations at three different interaction strengths at 250 mM salt. At ε=0.13 kcal/mol, a condition calibrated without crowders based on previous FRET experiments, we observe the slab’s complete disruption at 250 mM salt, aligning with experimental findings indicating no LLPS in the absence of crowders. At ε=0.15 kcal/mol, a small slab and additional small protein clusters emerge. Finally, at ε=0.17 kcal/mol, a fully formed slab is observed. To analyze these simulations and determine protein concentrations in low- and high-density phases, we computed the density profile along the z-axis (Fig. 2C). At ε=0.13 kcal/mol without crowders, stable slabs and LLPS do not exist for all salt conditions. Upon increasing ε to 0.15 kcal/mol, two well-defined phases coexist at 500- and 1000-mM NaCl. While a small slab is visible at 250 mM NaCl, quantifying coexistence densities may be challenging due to the disrupted slab. This simulation condition resembles the 20% PEG400 condition observed in the experiment (Fig. 2A). At ε=0.17 kcal/mol, LLPS occurs above 50 mM NaCl concentration, akin to the 25% PEG400 condition. This suggests that crowding-induced LLPS can be explained by heightened amino acid interactions resulting from increased volume exclusion under high crowding conditions.

To further investigate the salt dependence of LLPS, we calculated the protein concentrations in the low- and high-density phases by averaging the density profiles at the two ends and in the middle. As shown in Fig. 2D, we plot the concentrations of coexisting phases under various salt conditions. The concentration of the low-density phase is therefore the lowest concentration for the protein to phase separate, which is often referred to as the critical concentration. As salt concentrations increase under two distinct crowding conditions, the critical concentration decreases, indicating that LLPS can occur more readily at lower protein concentrations. Hence, salt induces LLPS, aligning with the experimental findings in Fig. 2A. The trend is monotonic, different from the turning behavior seen using the size of the protein in Fig. 1K, suggesting the correlation between the single-chain and LLPS behaviors also breaks down in the simulations. We further explore the intermolecular contact map between different chains which provides a better understanding of the LLPS behavior in contrast to the intramolecular contact map calculated using the single-chain simulation previously. We find there are clusters of contacts between NTD and CTT at low salt concentrations (Fig. S9), suggesting these specific interactions between NTD and CTT persist even across different chains. However, the pattern of these specific interactions starts to disappear when more uniform contacts are formed at high salt concentrations. To understand the salt dependence, we then plot the number of contacts between different regions of αSyn at different salt concentrations (Fig. S10). The contacts between different regions exhibit overall monotonic behavior, aligning with the monotonic salt dependent LLPS. This suggests that the LLPS behavior is influenced by the intermolecular interactions among all regions of αSyn, rather than being exclusively governed by the intramolecular NTD-CTT interactions.

### Decoupling aggregation and phase separation of αSyn

In both experiments and simulations, we previously observed the crowding and salt-induced LLPS of αSyn. Unlike the non-monotonic salt-dependent aggregation, the salt-dependent LLPS exhibits a monotonic trend. This can be attributed to intermolecular interactions between αSyn chains, in contrast to the aggregation salt dependence characterized by intramolecular NTD-CTT interactions. Simplifying the picture through coarse-graining along the αSyn sequence, the salt-dependent intermolecular interactions among αSyn chains are influenced by the overall net charge of αSyn (i.e., -9), leading to an overall net repulsive interaction between different chains. Increasing salt concentration and electrostatic screening can effectively reduce the repulsive interactions and then induce LLPS. On the other hand, aggregation is dependent on the intramolecular NTD and CTT interactions and the net charge within NTD (i.e. +4) and CTT (i.e.

-12) (Fig. 1A). αSyn, therefore, becomes a perfect example due to its special sequence arrangement, in which the sequence charges governing LLPS and aggregation differ. Salt becomes the easiest way to separate the sequence contribution to LLPS and aggregation since both can be understood by the interactions between charged amino acids.

We investigate further the salt dependent LLPS at a variety of protein concentrations (15 µM to 150 µM) in the presence of different PEG400 conditions. After 29 days of incubation along with 20% PEG, αSyn formed small droplets at high salt concentrations (250 to 1000 mM) at almost all protein concentrations (images highlighted in blue in Fig. 3A). However, at high protein concentrations (75 and 150 µM), macroscopic fibril-like structures could be observed (images highlighted in red). They can be up to a few hundred micrometers long. We would like to note for the four specific conditions: 75 µM protein concentration and 50 mM salt, and 150 µM protein concentration and three different salt concentrations (50, 75, 125 mM), there is no droplet, but fibril-like structure formation. At least in these conditions, this is different from what has been previously observed that αSyn nucleates from the liquid droplet(26). More specifically at another condition (150 µM protein concentration and 250 mM salt), we see the coexistence of both fibril-like structures and droplets, and they are not close to each other in the image provided. A simplified phase diagram can be found in Fig. 3B.

**Figure 3.**
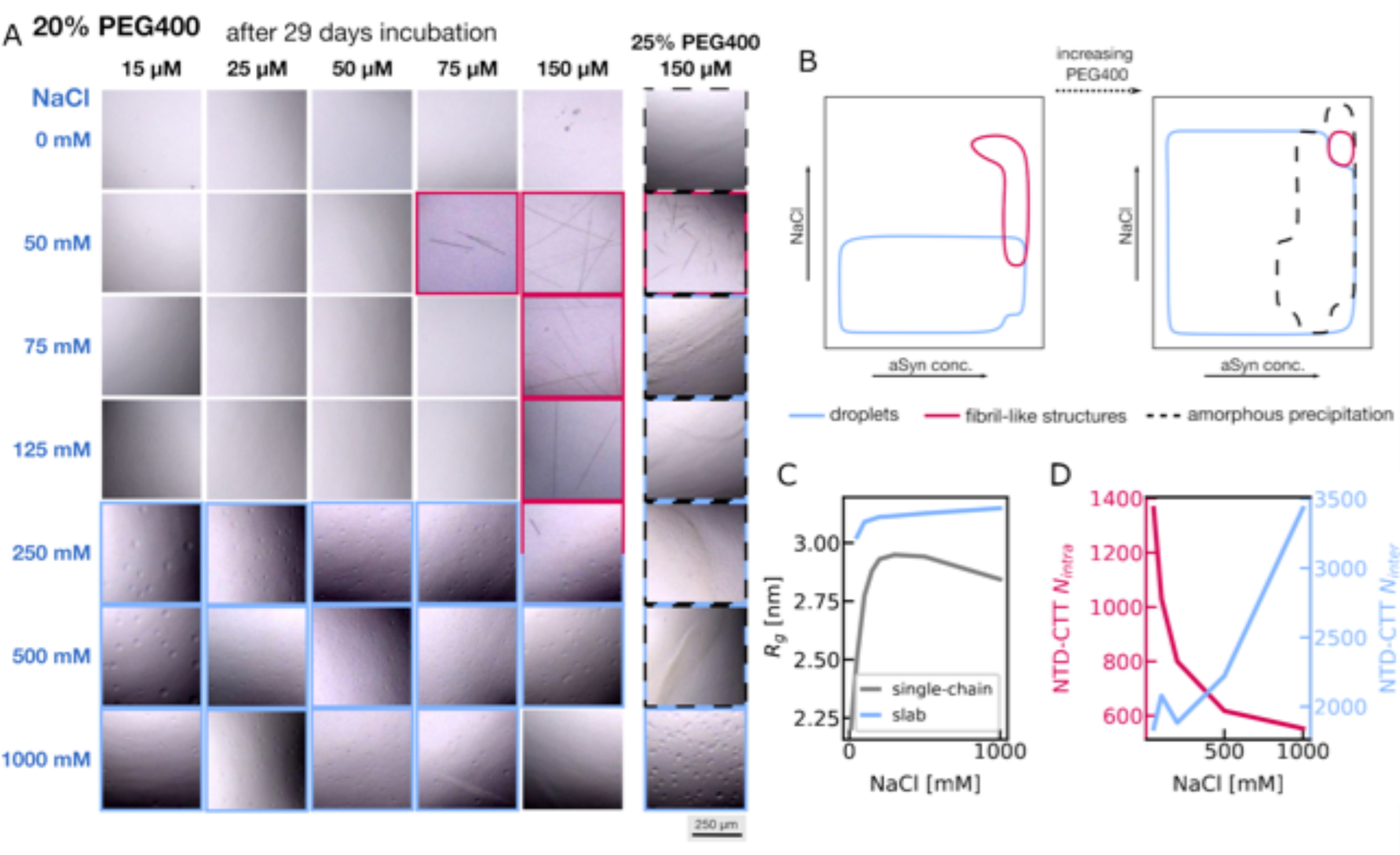
(A) Micrographs (500 x 500 µm) of LLPS formation of WT αSyn at pH 7 at different salt and protein concentrations after an incubation of 29 days in the presence of 20% PEG400. Dot plot at (B) 20% PEG400 and 25% PEG400. (C) The salt-dependent *R_g_* from the single-chain and slab simulations. (D) The intra- and intermolecular NTD-CTT interactions for αSyn in the slab simulation.

To confirm our observation that fibrils and droplets can emerge at different solvent conditions, we obtained the same set of images at 25% PEG (Fig. S11). At higher PEG concentration, we anticipate that αSyn will more readily form liquid droplets due to the excluded volume effect and increased interactions between amino acids. αSyn indeed formed droplets at all protein and salt concentrations (≥50 mM NaCl). At elevated protein concentrations, amorphous precipitation was observed, akin to observations during crystallization (black striped line) (Fig. 3A, S11). Unlike LLPS which occurs in the metastable zone, in a typical crystallization phase diagram, amorphous precipitations occur in a zone, which is too supersaturated to promote ordered aggregation(66). However, excluding amorphous precipitation the salt dependence of both aggregation and LLPS stays qualitatively the same as at 20% PEG: aggregation still occurs in medium salt concentration whereas LLPS occurs above the critical salt concentration. This is because the crowders have a more general impact on the interactions of between all types of amino acids instead of only charged amino acids. No macroscopic aggregation, precipitation, or droplet formation was observed after 29 days of incubation when the crowding agent was reduced to 18% (Fig. S12).

As shown previously, our coarse-grained simulation can explain the different salt dependences of aggregation and phase separation alone: aggregation is dependent on the intramolecular NTD-CTT interactions whereas LLPS is dependent on the overall intermolecular interactions between different chains of αSyn. However, due to the coexistence of droplets and aggregates at specific conditions, fibrils can possibly grow inside a droplet (Fig. S2). It is known from a recent experiment that αSyn adopts a more elongated conformation after LLPS(47). As shown in Fig. 3C, the *R_g_* values of αSyn were calculated from the single-chain simulation and from the slab simulation (ε=0.15 kcal/mol). αSyn becomes much more expanded under the LLPS condition, in agreement with the previous experimental observation. It would be interesting whether the intramolecular interactions before forming the liquid droplets stay with the same salt dependence as those after forming the liquid droplets. In Fig. 3D, the intramolecular NTD-CTT interactions in the slab simulation (ε=0.15 kcal/mol) decrease and then increase following the same trend as seen in a single-chain simulation. Surprisingly, we now see completely opposite salt dependence for the intra- and intermolecular interactions of αSyn coming from the same simulation under the LLPS condition. This is consistent with both the salt dependence of aggregation and phase separation observed from the experiment.

To further illustrate the sequence features resulting in the opposite salt-dependent intra- and intermolecular interactions, we examined two variants of αSyn through coarse-grained simulations. We hypothesized that such an opposite dependence is due to the unbalanced net charges in different regions of αSyn: NTD (+4), NAC (-1) and CTT (-12). For the first variant, we mutated nine of the negatively charged amino acids in CTT to polar amino acids with similar sizes so that now the net charge of the entire sequence is zero (Fig. S13A). Electrostatic screening now has limited contribution to its intermolecular interactions and consequently salt dependent LLPS at low salt concentrations below 300mM (Fig. S13B). LLPS slightly favors high salt concentrations above 300mM due to the salting-out effect (56). Yet, there displays a turning behavior of *R_g_* undervaried salt concentrations because of the slight attraction between NTD and CTT (Fig. S11D). However, both intra- and intermolecular interactions now exhibit the same salt dependence (Fig. S13B), suggesting the absence of the heteropolymeric behavior which was observed in wild type αSyn shown in Fig. 3D. We further introduce a second variant in which the net charge in each of the three regions of αSyn is fully balanced. For NTD, the unbalanced charges all come from the four EKTK or KTKE segments, which has been previous targeted to disrupt fibrillation(67). Here we mutated one of the Lys in each segment, so that the local net charge is zero (Fig. S13A). To balance NAC and CTD, we also mutated a certain number of negatively charged amino acids to polar ones. This variant is expected to behave completely like a homopolymer. We now observe monotonic salt dependence for all parameters calculated, including intra- and intermolecular interactions, density of the droplet and *R_g_* (Fig. S13B, C and D). This verifies our hypothesis that, the opposite salt dependence in intra- and intermolecular interactions, and consequently the different salt dependence in LLPS and aggregation, are attributed to the unbalanced arrangement of the charged amino acids in αSyn.

### Dissolution of αSyn aggregates into droplets

Previous work observed that αSyn aggregate nucleates from its droplet state(26). Here, we observed that phase separation and aggregation have completely independent salt dependence. This might not be contradictory if the two states possess different free energies with a high kinetic barrier separating them supported by a most recent work(68). Then the final state is solely dependent on the free energies of the two states at a specific solvent condition, whereas the first appeared state might only be due to its faster kinetics and could differ from the equilibrated state. When observing at different solvent conditions, one might see droplet or fibril alone, or coexistence of fibril and droplet as what we observed (Fig. 3). When observing at different time windows, one might see the interconversion between the droplet and aggregate states depending. We therefore investigate the droplet and fibril formation at different incubation time.

Strikingly after 120 days long incubation, αSyn only forms droplets under all investigated conditions: at different salt and protein concentrations and at different crowding (18, 20 and 25%) and pH (pH 7 and 6.2) conditions (Figure 4A, B and C). The fibril-like structures observed at high protein and low salt concentrations after 29 days of incubation have all disappeared (see Fig. S11, S12, S14 and S15). This implies that the droplet state has a lower free energy at these conditions, making it more stable. However, the amorphous aggregates formed at stronger crowding conditions appeared unchanged during the incubation time, suggesting that these assemblies are more stable than both fibril and droplet. The long incubation time also points out the high kinetic barrier of dissolving the aggregates, making this type of experiment extremely difficult to achieve equilibrium. The 120 days incubation might still not be sufficiently long, and the incubation time is also dependent on the solvent conditions. To confirm that the disappearance of aggregates is not due to protein degradation, we have performed mass spectrometry on a sample containing droplets after initial aggregation and subsequent incubation for 140 days Fig. S16). We further verified the liquid-like properties of droplets by dilution and droplets disappeared within 90 seconds after dilution (Fig. S17).

**Figure 4.**
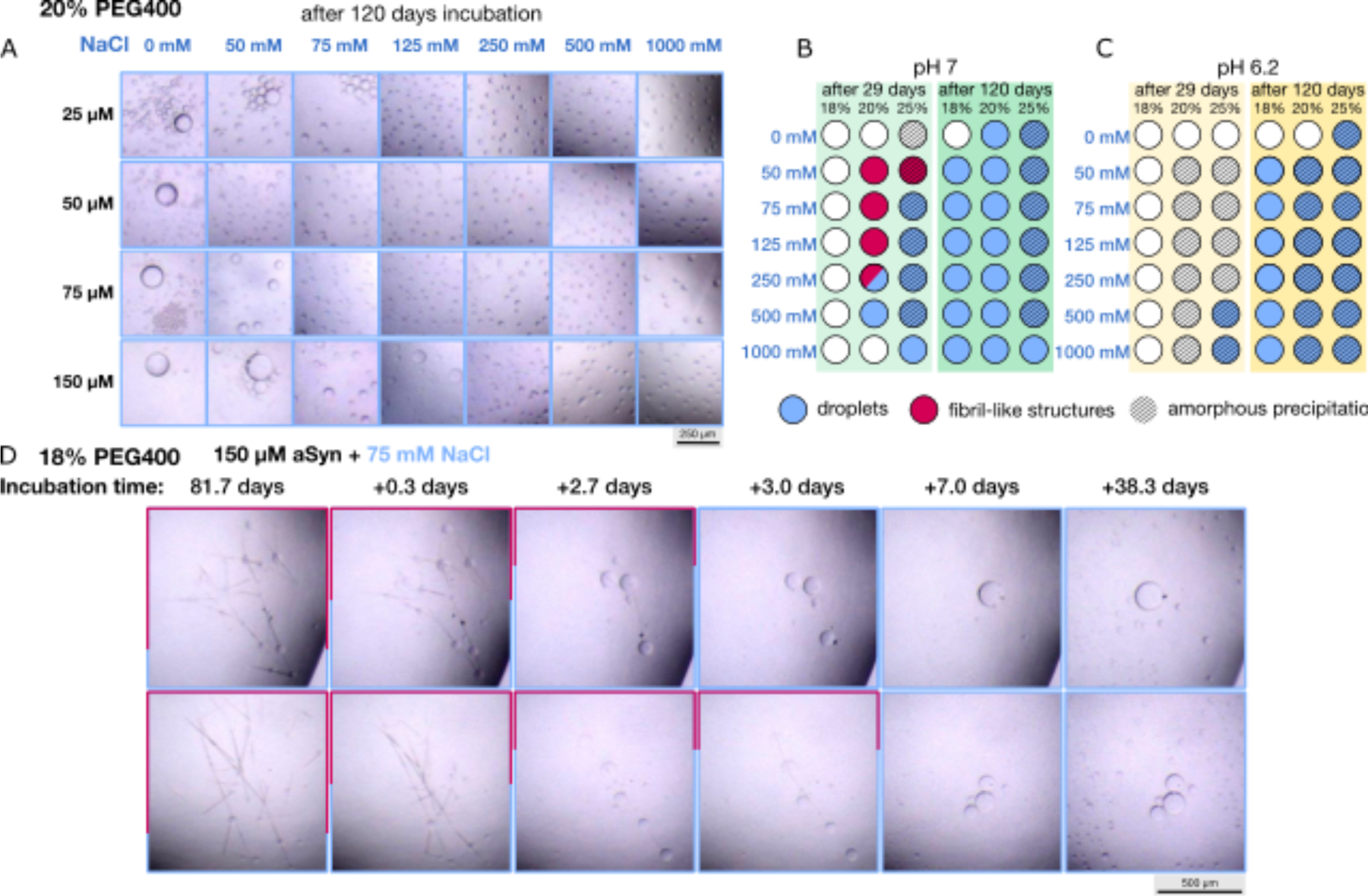
(A) Micrographs of droplet formation of WT αSyn at pH 7 at different salt and protein concentrations after an incubation period of 120 days in the presence of 20% PEG400. Dot blot at pH 7 (B) and pH 6.2 (C) of 150 µM αSyn in the presence of 18%, 20% and 25% PEG400. The status after 29 days of incubation is displayed on the left (light color) and after 120 days of incubation on the right (dark color). The presence of droplets is indicated with blue, fibrillar structures in red and amorphous precipitation with stripes. (D) Exemplary micrographs of 150 µM αSyn in the presence of 18% PEG400 at 75 mM salt (bottom) over time. Fibrillar structures are dissolved by droplets.

To capture how these fibrils convert to droplet on the fly, we select a lower PEG concentration of 18% with the motivation that the diffusion and interconversion might be faster than these at 20% PEG. We prepare two identical sample of 150 µM WT αSyn at 18% PEG and 75 mM salt, and start imaging once every one or two days, trying to capture the exact time window when the fibril starts to covert to droplet. As shown in Fig. 4D, for both samples, macroscopic fibril-like structures formed commenced rapidly within a few days and persisted until day 81. At the 81^st^ day, we started seeing droplets growing and coexisted with fibril-like structures. Then with slightly different kinetics for the two samples, fibril-like structures seem to be dissolved by the droplets and completely disappeared at about 86^th^ to 89^th^ day. Looking more specifically the droplet position to dissolve the fibril, we find that these droplets with the ability of dissolving the fibrils are usually at the intersection points of two fibrils or on top of one fibril. This provides strong evidence that the droplet and aggregate states are two metastable states with a kinetic barrier between them. Their interconversion can be triggered when droplet starts developing close to the aggregate or when fibril starts to grow inside the droplet as observed by others. Whether we see the fibril or droplet depends on the free energy of the two assembly states at a specific solvent condition. If the incubation time is sufficiently long, then the appearance of the assembly state is also dependent on the kinetics.

Given the strong dependence of observation time on the assembly state of αSyn, we further investigate the dependence of the size of the droplet and the length of the fibril-like structures to their kinetics. While the droplets demonstrated a salt-dependent size distribution after 29 days of incubation with a decrease in size with increasing salt and protein concentration, the size distribution after 120 days of incubation is very similar throughout the different salt and protein concentrations (Fig. 5A). The largest droplets can be observed at a medium salt concentration of 75 mM. In the absence of salt, αSyn formed a single large phase over 100 µm in size. All protein concentrations demonstrated the fastest kinetics at 500 mM salt (Fig. 5B). The fastest droplet formation could be observed at 150 µM αSyn in the presence of 500 mM salt, but was slowed down massively at 75 µM protein concentration. Given the noise of these data sets, we were not able to obtain a clear correlation between the size of the droplet and the kinetics.

**Figure 5.**
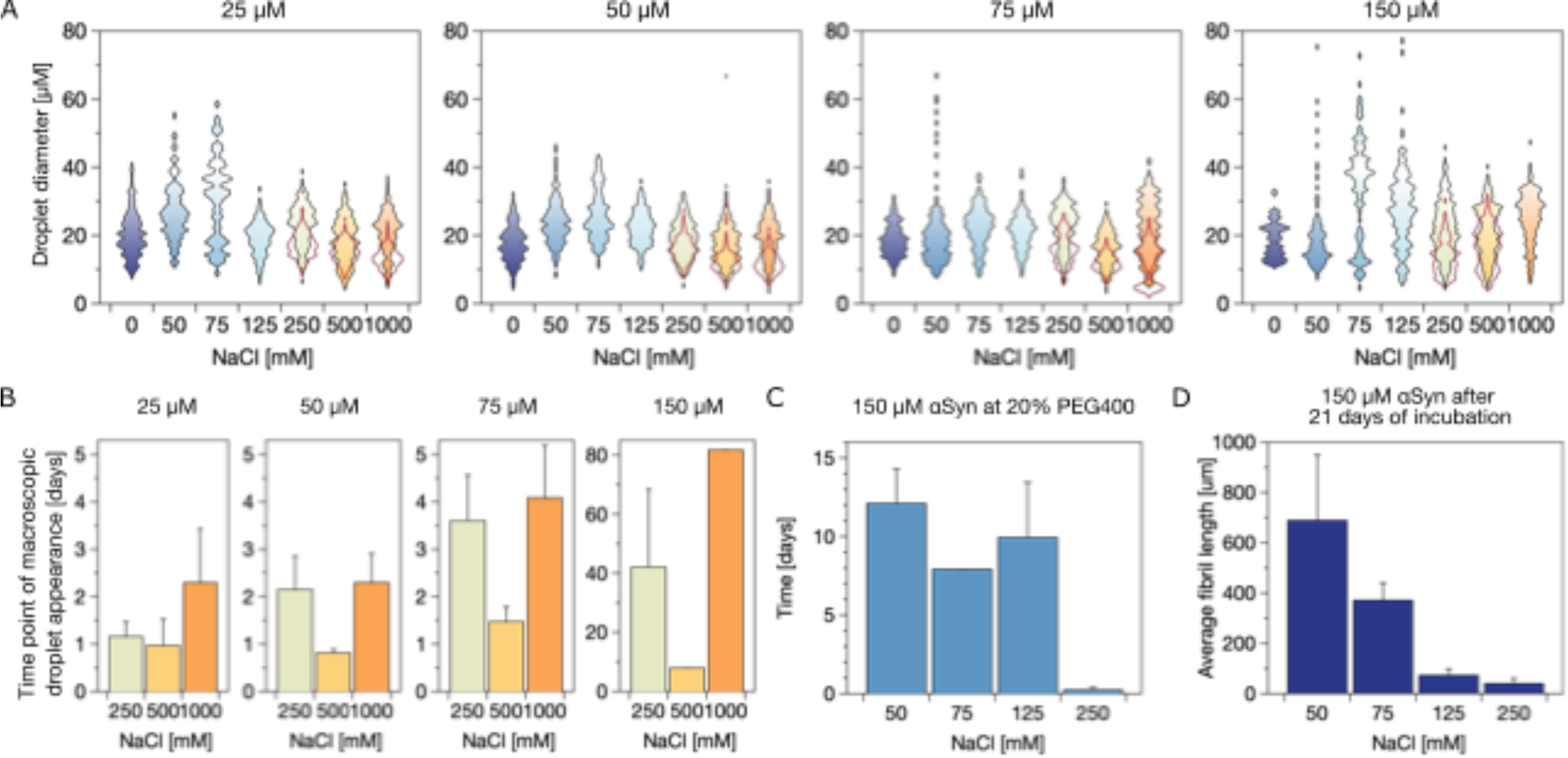
(A) Size distribution of droplets at different WT αSyn concentrations (25 µM, 50 µM, 75 µM and 150 µM) after an incubation period of 120 days in the presence of 20% PEG400 at pH 7. The size distributions after 29 days at high salt concentrations (250 mM, 500 mM, and 1000 mM) are marked with a red line. The single large phase at low salt concentration is not displayed. (B) Average time point of appearance of macroscopic droplets at high salt concentrations. (C) Time point of appearance macroscopic fibrillar bundles of 150 µM αSyn at 20% PEG400 and (D) their length after 21 days of incubation.

On the other hand, we can estimate the size of the fibril and compare that with the time of observing the fibril. The correlation between the size and kinetics of fibril is better interpreted in contrast to those of the droplet. As shown in Fig. 5C, macroscopic fibril-like structures of 150 µM αSyn appear later at low salt concentrations, though with a longer average length after 21 days of incubation. Fibrils are also thinner and more bent at low salt concentrations. We noted a distinct correlation between the size of the fibril and its kinetics, wherein slower growth corresponds to larger assembly. As salt concentrations increase, reducing interactions between NTD and CTT, the fibril initially appears to grow faster due to reduced hindrance from CTT in NTD aggregation. However, the increasing intermolecular NTD interactions may introduce more kinetically arrested structures, slowing down growth as a longer time is needed for structural rearrangement. A similar trend was observed in a previous study on droplet growth (69), where stronger interactions led to many small droplets, kinetically trapped in an arrested state instead of growing into one large droplet.

## Discussion and Conclusion

Both phase separation and aggregation of αSyn including the interplay between the two assembly states have gained tremendous interest recently. In previous studies, altering solvent conditions or introducing different variants of αSyn often had similar effects, promoting or inhibiting both aggregation and phase separation. Here, we explore the phase separation and aggregation of αSyn by systematically varying protein, salt, and crowding concentrations at multiple pH levels using two different variants of αSyn. This approach enables us to observe the distinct behaviors of phase separation and aggregation independently.

Specifically, the salt dependence of aggregation exhibits a non-monotonic pattern, with aggregation primarily occurring in medium salt concentrations shown in Fig. 6A (left panel). This is mainly caused by the modulation of intramolecular NTD-CTT interactions through the interplay of electrostatic screening and the salting-out effect. CTT and its interaction with NTD were previously known to inhibit αSyn aggregation(14). Displayed in Fig. 6A (right panel), the salt dependence of LLPS is monotonic with phase separation initiating as salt concentration increase. This is attributed to the fact that phase separation depends on the intermolecular interactions among all regions of αSyn rather than solely relying on either the NTD or the CTT. Therefore, the sequence determinant of aggregation is the net positive charge of NTD (+4) and net negative charge of CTT (-12), whereas that of phase separation is the overall net charge of the entire chain (-9). The unique sequence arrangement of αSyn and its specific heteropolymeric interactions between NTD and CTT contribute to the decoupling of its single-chain and liquid-liquid phase separation (LLPS) behaviors. This decoupling is distinctive compared to many other intrinsically disordered proteins where these behaviors are often assumed to be correlated. In addition, one of the most notable findings in our studies is that the assembly state of αSyn is dependent on the duration of incubation time. Specifically, at high salt concentrations exceeding 250 mM, only macroscopic LLPS is observed for all tested conditions (Fig. 6B right). However, at low salt concentrations below 250 mM, various scenarios unfold, including macroscopic fibril-like structures only, droplet only, and the coexistence of fibril and droplet under different protein and crowding conditions (Fig. 6B left). Fibril forms early in our solvent condition characterized by close-to-neutral pH and low salt concentration, but eventually all convert into droplets after prolonged incubation. This observation challenges common assumptions about the stability of fibril structures, which are typically considered resistant to dissolution without altering conditions such as solutions, ligands, or post-translational modifications. However, by taking images every one or two days after 80 days of incubation, we were able to capture the gradual conversion of αSyn fibril into droplet in real time. The droplet formation frequently occurs at the intersection of two fibrils or at the top of a single fibril, ultimately leading to the complete dissolution of the fibril. It is crucial to highlight that our study is not contradictory to previous observations regarding fibril growth inside droplets. Due to the distinct sequence determinants of the droplet and aggregation states, these two states can be operated independently of each other when varying the solution conditions or protein concentrations. Therefore, whether aggregates nucleate inside droplets or aggregates convert into droplets depends on specific experimental conditions. Only by systematically testing various salt, protein, and crowder conditions, as done in this study, can all possible scenarios be explored. Owing to technical constraints in our fibrillation and LLPS assays, we were unable to carry out these experiments under identical conditions. It is crucial to emphasize that we upheld uniformity in buffer composition and protein conditions for both assays. This strategy allows for a systematic comparison of the effects of salt concentration, crowding, and pH on LLPS and fibrillation.

**Figure 6.**
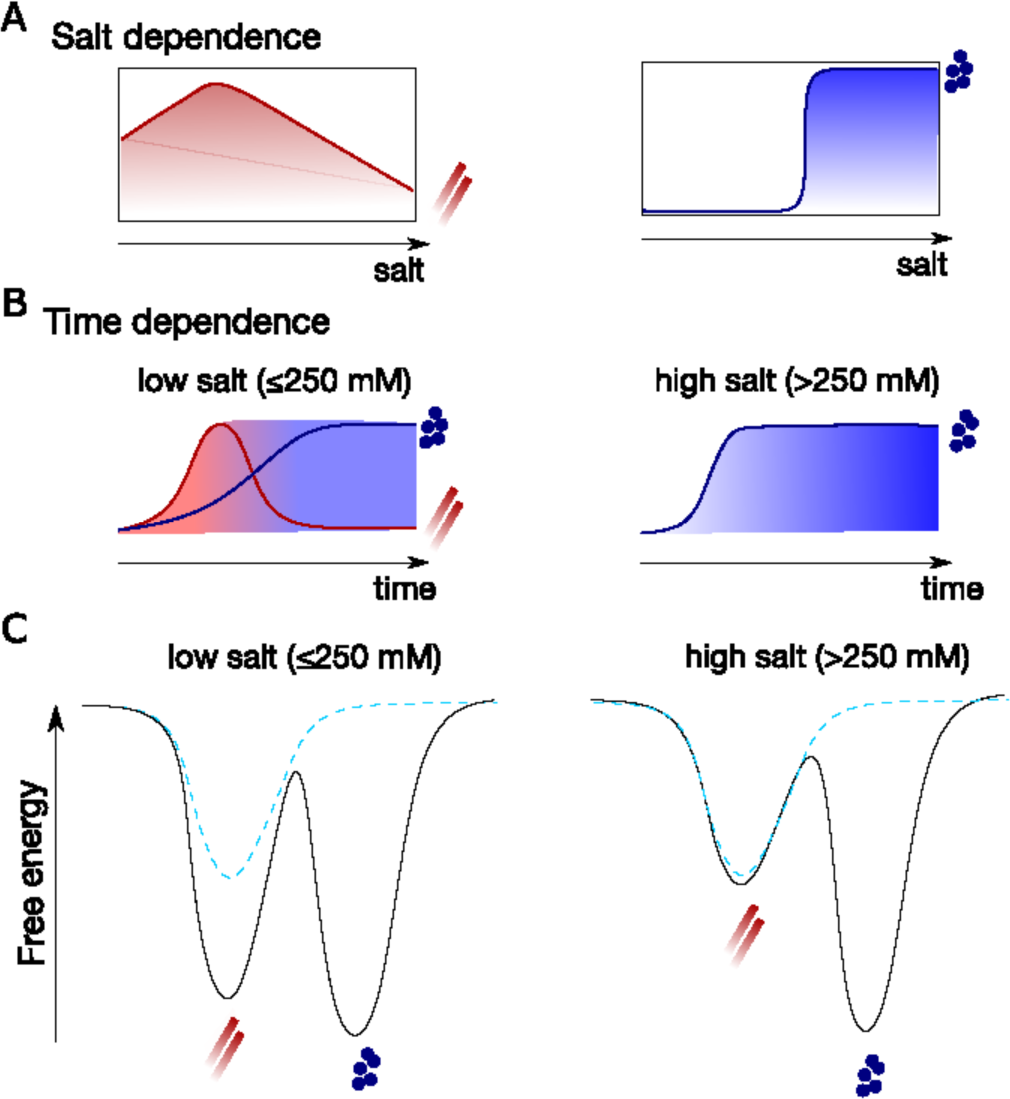
Schematic diagram of the interplay between the αSyn solid aggregates and liquid droplets. (A) Salt-dependent aggregation and phase separation. (B) Time-dependent aggregation and phase separation. (C) A free energy landscape includes the two assembly states at crowding conditions (solid line) and without crowders (dash line).

Then how can we understand such a counterintuitive observation in which fibrils can be dissolved by droplets? Polymorphism is often observed for many of these fibril-forming proteins including αSyn, in which there can be a variety of fibril structures depending on the conditions(70, 71). There have also been proposed that the formation of different polymorphic fibril structures is under kinetic control(72) and can be described by an energy landscape(73). Motivated by these previous studies, we propose an energy landscape picture where αSyn aggregation and phase separation occur at different crowding and salt conditions. Without crowders at all salt concentrations, the only stable assembly state of αSyn is aggregate (dash lines in Fig. 6C). As crowding conditions increase, both the aggregate and droplet states become more stable. Under conditions of low salt concentration (as shown in Fig. 6C on the left), the formation of aggregates was observed to precede the formation of droplets. This observation may be attributed to the interplay between percolation and phase separation, as well as the network structure within the αSyn droplet (63, 74), necessitating additional investigation. However, the droplet state is more stable and at last, all aggregates slowly convert into droplets due to the significant kinetic barrier between the two states. At high salt concentration (Fig. 6C right), the droplet exhibits increased stability, while the aggregate shows reduced stability owing to their distinct salt dependence, and therefore only macroscopic droplets were observed. Nonetheless we cannot exclude whether there exist structural features near the droplet surface, which might disrupt instead of promoting fibril-like structures (75).

The commonly observed correlations between phase separation and aggregation in proteins that exhibit both behaviors are likely attributed to the escalating local protein concentration and the more expanded conformation of the protein. Both factors contribute to the conversion of liquid droplets into solid aggregates. However, our findings, at least in the case of αSyn, may reshape our understanding of the interplay between its LLPS and aggregation. The sequence features of the two assembly states may not necessarily co-evolved given the different functional and/or pathological relevance of the two states. Additionally, the sequence determinant of LLPS and aggregation is also not necessarily the same due to the intrinsic physics governing the formation these two assemblies. A meticulous investigation under various solvent and protein conditions might offer a different perspective on the interplay between LLPS and aggregation for some other proteins. The droplet state could be potentially functional relevant to inhibit aggregation as a recent literature suggested(51), or even dissolve the aggregate through a variety of heterotypic interactions saving the protein from its dynamically arrested state(76).

## Material and Methods

### Protein sample preparation

WT αSyn1-140 and ΔCTT variant αSyn1-108 were expressed in *E. Coli* BL21(D3) and purified as described previously. Briefly, the expression was conducted for 24 h at 20°C in LB-medium and harvested. The protein was precipitated using ammonium sulphate after lysis of the cell pellet by heat and ultrasonication. The protein was re-suspended in 25 mM Tris-HCl, pH 8, filtered (0.42 pore size) and subsequently loaded onto a HiTrapQHP anion exchange chromatography column (GE Healthcare). The protein was eluted using a gradient with 25 mM Tris-HCl, pH 8, 800 mM NaCl and the αSyn containing fractions were combined, precipitated with ammonium sulphate and the pellet was stored at -20°C. The pellet was re-suspended and size exclusion chromatography was performed using a Superdex 200 Increase 10/300 GL. αSyn concentration was determined by measuring UV-absorption at 274 nm (extinction coefficient of 5600 M-1cm-1), aliquoted and lyophilised after dialysis against ammonium bicarbonate.

### Measurement of aggregation kinetics

The salt dependence of aggregation was investigated by evaluating ThT-aggregation kinetics performed in the presence of different NaCl concentrations (final concentrations: 0 mM, 10 mM, 50 mM, 75 mM, 125 mM, 250 mM, 500 mM, 1000 mM, 1500 mM, 2000 mM). Protein solutions were diluted to a final concentration of 15 µM. The aggregation kinetics were observed under crowding conditions (20% PEG400) and in the absence of a crowding agent in 30 mM phosphate buffer with a final ThT concentration of 20 µM.

Three technical replicates of each solution were then pipetted (35 µl each) into a 384er non treated PS surface plate (Thermofisher). Before sealing the plates using aluminium adhesive foil (neoLab), a small glass bead (diameter 1.0 mm) (Sigma) was added to each well. The kinetics of amyloid fibril formation were monitored at 37°C under continuous shaking (300 rpm) conditions by measuring ThT fluorescence intensity through the bottom of the plate using a PHERAstar (BMG LABTECH, Germany) microplate reader (readings were taken every 150 seconds). In addition, single monomer concentrations were tested with shaking or under quiescent conditions.

In order to compare the factor of increase of the ThT fluorescence emission intensity between the samples, the maximum ThT fluorescence emission intensity was compared with the lowest emission value. The halftimes of the aggregation reaction are defined as the point where the ThT intensity is halfway between the initial baseline and the final plateau. The halftimes were obtained by individually fitting the curves using a generic sigmoidal equation (nielsen2001).

Single conditions were chosen to analyze the morphology of formed aggregates with atomic force microscopy. The samples were diluted to a final concentration of 10 µM and 10 µl were applied to a freshly cleaved mica surface, and incubated for 5 min. The samples were washed 5 times using 100 µl dH_2_O and subsequently dried under a gentle flow of N2. The samples were investigated using a Dimension Icon (Bruker) in tapping mode.

### Measurement of droplet formation

The LLPS experiments were conducted in MRC3 crystallization plates. The samples were prepared by using the multi-channel pipetting robot mosquito (SPT labtech), 500 nl protein was mixed 1:1 with the buffer solution to achieve different salt (0 mM, 50 mM, 75 mM, 125 mM, 250 mM, 500 mM, 1000 mM) and crowding agent (18%, 20%, 25%) concentrations. At least 4 replicated per condition were investigated. The plates were sealed and imaged using a Rock Imager microscope (Formulatrix). The plates were kept at 20°C and imaged after certain time points. The droplet size and fibril length were assessed using Fiji.

### ESI-TOF LC-MS analysis after long incubation

50 µM aSyn sample in the presence of 75 mM NaCl and 20% PEG400, which was aggregated for 62 h at 37°C and subsequently incubated at RT for 140 days, was analysed using the Waters LTC Premier ESI-TOF mass spectrometer coupled with the Waters 2795 Alliance HPLC for separation. Before injection, GdnHCl was added to the sample to a final concentration of 1 M and incubated for one hour at RT. The MassPREP Micro Desalting Column was employed with a specific gradient to separate a maximum amount of detergent before the protein enters the mass spectrometer (MS). Analysis and deconvolution was made with the software MassLynx4.1 and MaxEnt1agorithm (Waters, Milford, USA).

### SAXS measurement

Synchrotron SAXS (small angle X-ray scattering) experiment was performed at the ID02 beam line at the European Synchrotron Radiation Facility (ESRF) (Grenoble, France). Sample solutions with 150 µM monomer concentration of αSyn and varied salt concentrations were measured at 20°C using a liquid sample changer coupled to a flow-through capillary cell (quartz, inner diameter 2 mm). 10 positions along the capillary length were recorded with a exposure time of 0.03 s. Background measurements with the corresponding buffer were taken before each sample using the identical positions, and measurement protocol (77). The datasets were processed with PRIMUS (78). The radius of gyration was estimated using Guinier analysis (79) as detailed in Fig. S5 and S6.

### Molecular dynamics simulations

We applied a previously developed residue-based coarse-grained model, HPS model (55), which was parameterized for studying liquid-liquid phase separation of intrinsically disordered proteins (IDPs) (61). In the HPS model, each amino acid was represented by a bead with its charge and hydropathy (80). There were three types of interactions: bonded interactions, electrostatic interactions and short-range pairwise interactions. The bonded interactions were characterized by a harmonic potential with a spring constant of 10 kJ/Å^2^ and a bond length of 3.8 Å. The electrostatic interactions were modeled using a Coulombic term with Debye-Hückel electrostatic screening(81) to account for the salt concentration. The short-range pairwise potential accounted for both protein-protein and protein-solvent interactions with an adjustable parameter for the interaction strength, which can be optimized using the experimental measurement of the size of α-Syn. We further added additional terms for angle and dihedral preferences: a statistical angle potential from a previous literature(82) for all types of amino acids and a statistical dihedral potential published previously(83). We found an interaction strength ε of 0.13 kcal/mol for the short-range pairwise potential best captured the FRET measurement of α-Syn(57), which is the same as what we found for Aβ40 in a previous work(58).

In order to simulate α-Syn at a low pH, we calculated the averaging charge of D and E based on their pKa values and changed the charge of these two amino acids in the CG model accordingly. For example, the pKa values of D and E are 3.65 and 4.25, respectively. At pH 3, the averaging charge of D and E are therefore -0.183 and -0.053, respectively.

The HOOMD-Blue software v2.9.3 (84) together with the azplugins (https://github.com/ <u>mphowardlab/azplugins</u>) were used for running the molecular dynamics simulations. All simulations were run using a Langevin thermostat with a friction coefficient of 0.01ps^-1^, a time step of 10 fs and a temperature of 298K. For single-chain simulations, the simulations were run for 5µs and the first 0.5µs were dumped for equilibration. For co-existence simulation using a slab initial conformation, the simulations were run for 5µs and the first 1µs were dumped for equilibration. The analysis was done using MDAnalysis (85).

## Acknowledgments

We acknowledge the support from the Swiss National Scientific Foundation (310030_197626, J.L.), and the Brightfocus foundation (A20201759S, J.L.), the National Science Foundation (MCB-2015030, W.Z.), the National Institutes of Health (R35GM146814, W.Z.), and the research computing facility at Arizona State University (W.Z.).

## Author Contributions

R.S-H, W.Z., and J.L. designed research; R.S-H, X.S., A.M., M.W., W.H., W.Z. and J.L. performed research; R.S-H., W.Z. and J.L. analyzed data; and R.S-H., W.Z., and J.L. wrote the paper.

