## Supplementary material for "Phase Separation and Aggregation of α-Synuclein Diverge at Different Salt Conditions"

### Supporting Figures

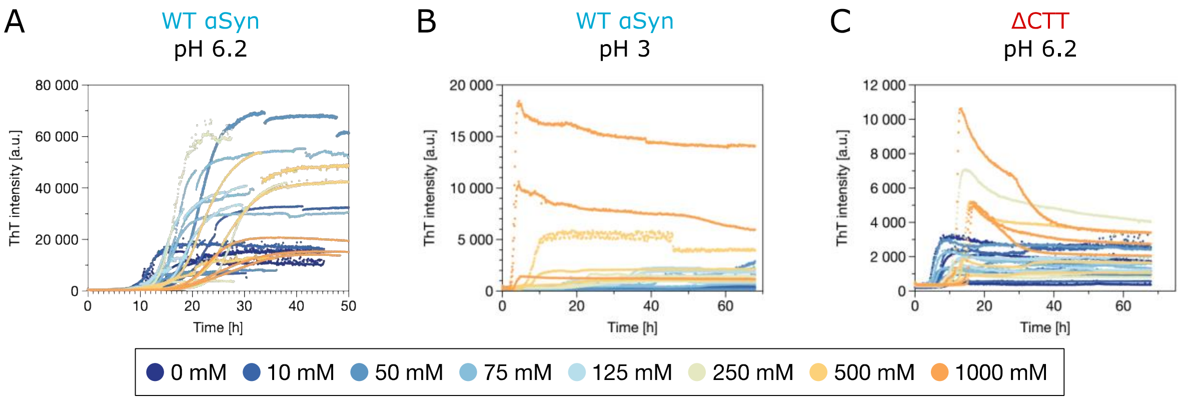


**Fig. S1**: Raw ThT fluorescence aggregation kinetics (see figure 1) at crowding conditions (20% PEG400), 37°C and under shaking conditions of WT α-Syn at pH 6.2 (A), pH 3 (B) and ΔCTT at pH 6.2 (C).

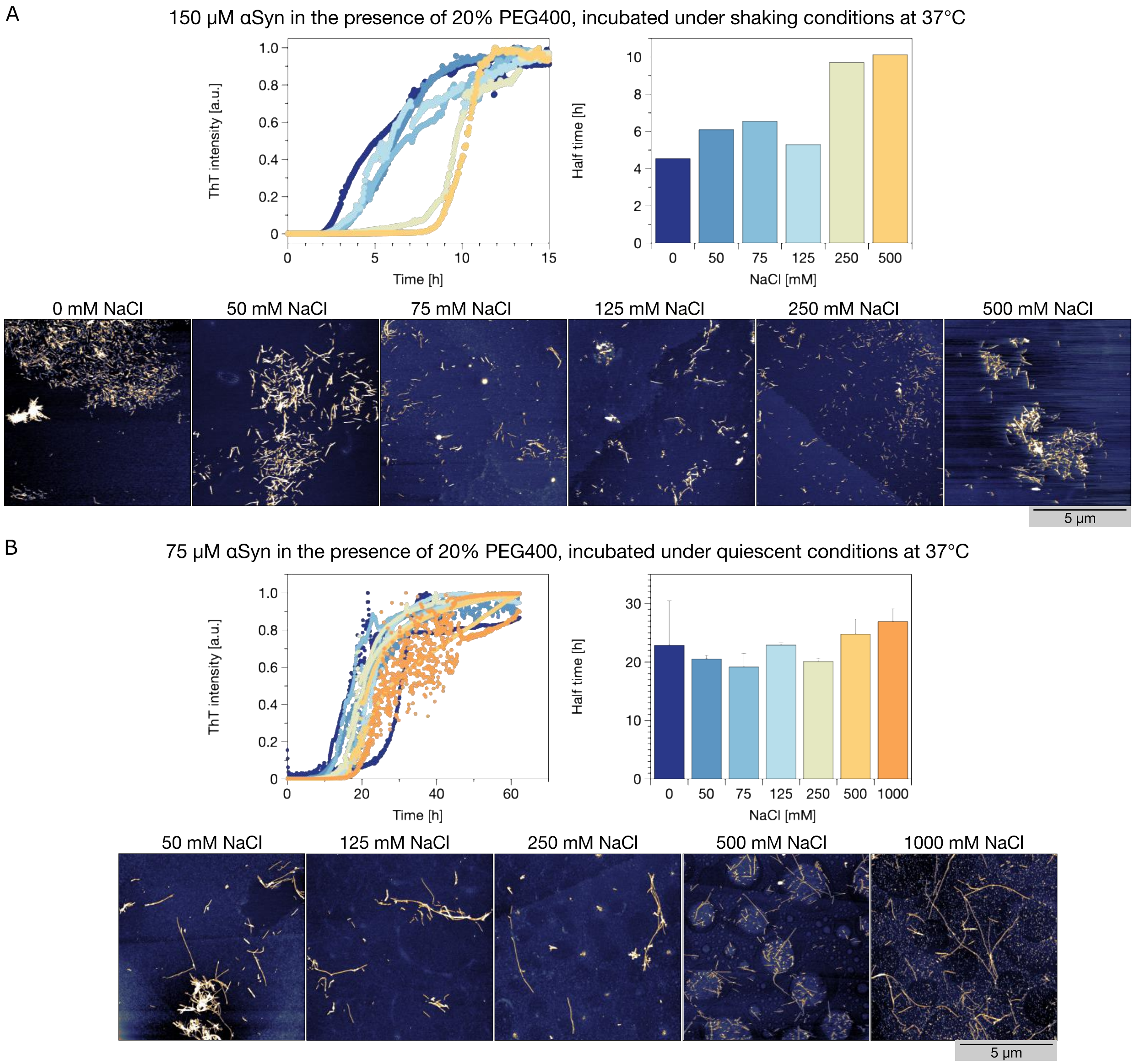


**Fig. S2**: Salt-dependent aggregation at pH 7 under crowding conditions (20% PEG400), 37°C of (A) 150 µM under shaking conditions (1 replicate) and (B) 75 µM WT αSyn under quiescent conditions (3 replicates). Normalized ThT-fluorescence aggregation kinetics and the half time of aggregation are displayed. A sample from the plateau-phase was taken and analyzed using atomic force microscopy. AFM imaging revealed the presence of amyloid fibrils at all investigated salt concentrations.

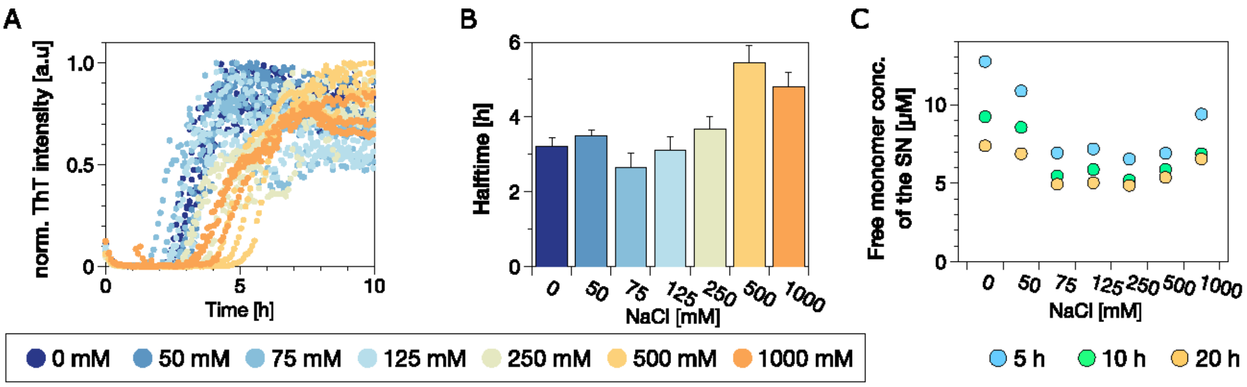


Fig. S3: A) Salt dependent aggregation of 15 µM WT αSyn in the presence of different salt concentrations under crowding (20% PEG400) and shaking conditions at 37°C (B) with the corresponding half times. C) After certain time-points (5 h, 10 h and 20 h) the samples were centrifuged (17 k xg) for 90 min and the free-monomer concentration of the supernatant was determined.

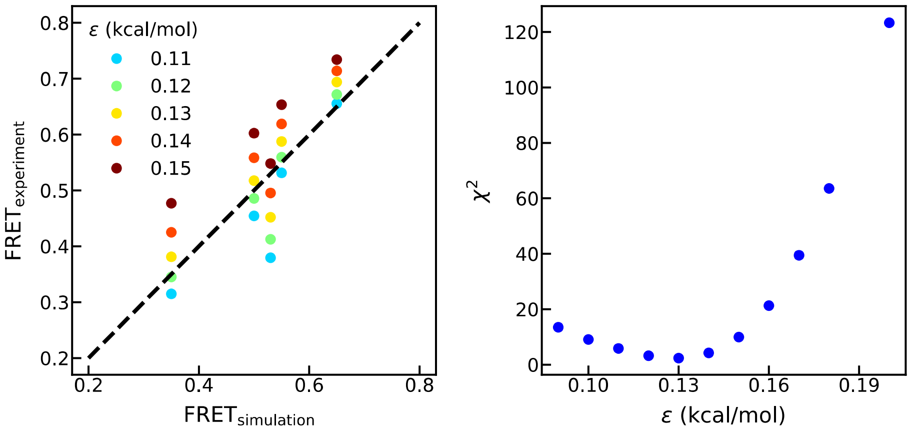


**Fig. S4**: The optimization of the interaction strength ε using the existing FRET data with five pair labeling positions [Nath *et. al*. Biophys. J. 103, 1940–1949 (2012)]. Left: Comparison between FRET efficiency from the experiment and from simulations with different interaction strengths shown in the legend. Right: The deviation of the simulated FRET efficiencies from experimental ones as a function of the interaction strength. ε=0.13 kcal/mol is the best to match the FRET experiment without crowding.

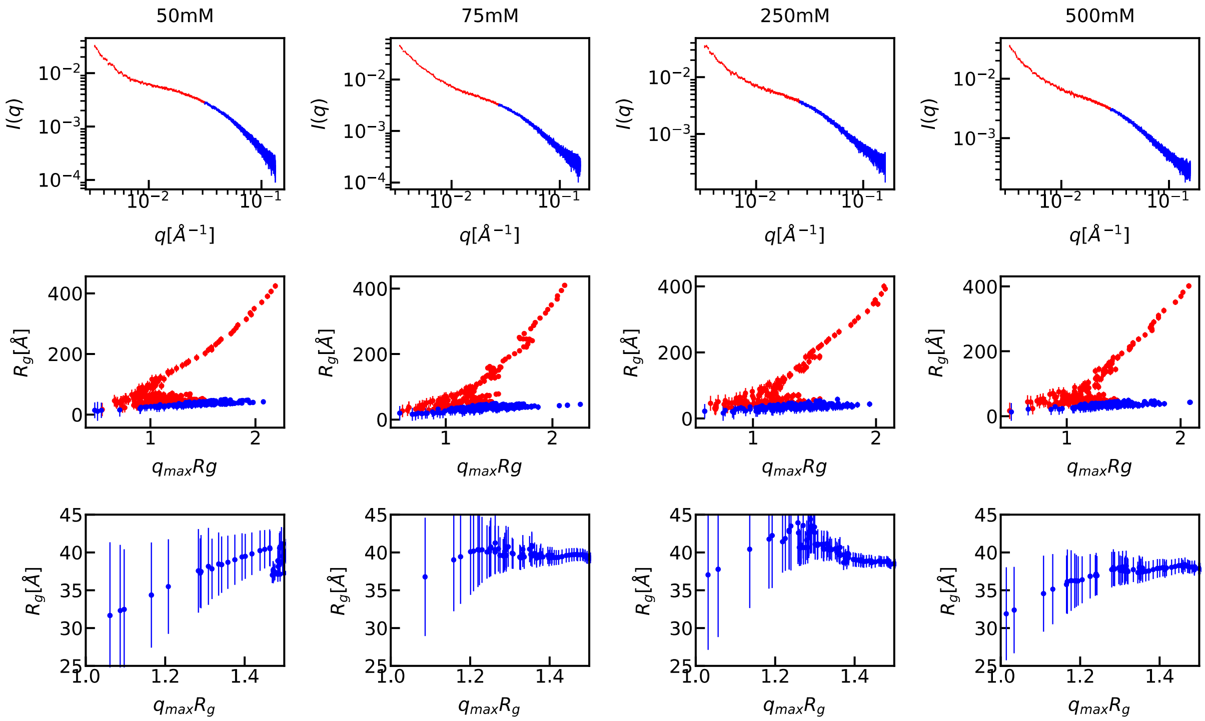


**Fig. S5:** Top row: SAXS intensities for αSyn at various salt concentrations, as indicated in the titles. Given the presence of particles with diverse sizes in the solution, we determined the radius of gyration ($R_g$) through Guinier analysis, employing a sliding window of 15 data points and a scattering vector q range of approximately 0.002 Å$^{-1}$. Medium row: The obtained $Rg$ plotted against the product of the maximum scattering vector $q_{max}$ for fitting and $R_g$. Two distinct regimes (highlighted in red and blue) with a turning behavior in between were observed, suggesting the prevalence of two different particle sizes. The first regime corresponds to aggregates or oligomers, while the second is expected to represent monomers. Using the turning point $q$ value, the SAXS intensities were partitioned into two regions (colored in red and blue in the top row). Bottom row: $R_g$ depicted as a function of the product of the maximum scattering vector $q_{max}$ for fitting and $R_g$, utilizing only the blue region of the SAXS intensities.

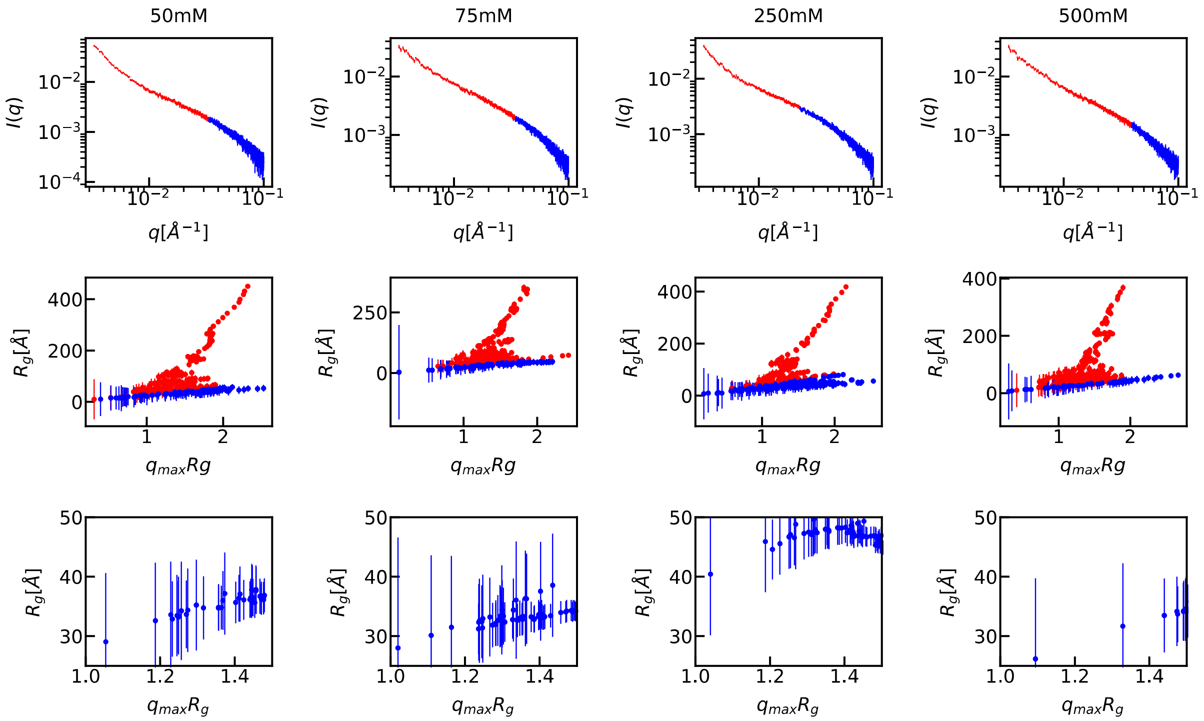


**Fig. S6:** Top row: SAXS intensities for αSyn at 20% PEG400 and various salt concentrations, as indicated in the titles. Given the presence of particles with diverse sizes in the solution, we determined the radius of gyration ($R_g$) through Guinier analysis, employing a sliding window of 15 data points and a scattering vector q range of approximately 0.002 $Å^{-1}$. Medium row: The obtained $Rg$ plotted against the product of the maximum scattering vector $q_{max}$ for fitting and $R_g$. Two distinct regimes (highlighted in red and blue) with a turning behavior in between were observed, suggesting the prevalence of two different particle sizes. The first regime corresponds to aggregates or oligomers, while the second is expected to represent monomers. Using the turning point $q$ value, the SAXS intensities were partitioned into two regions (colored in red and blue in the top row). Bottom row: $R_g$ depicted as a function of the product of the maximum scattering vector $q_{max}$ for fitting and $R_g$, utilizing only the blue region of the SAXS intensities.

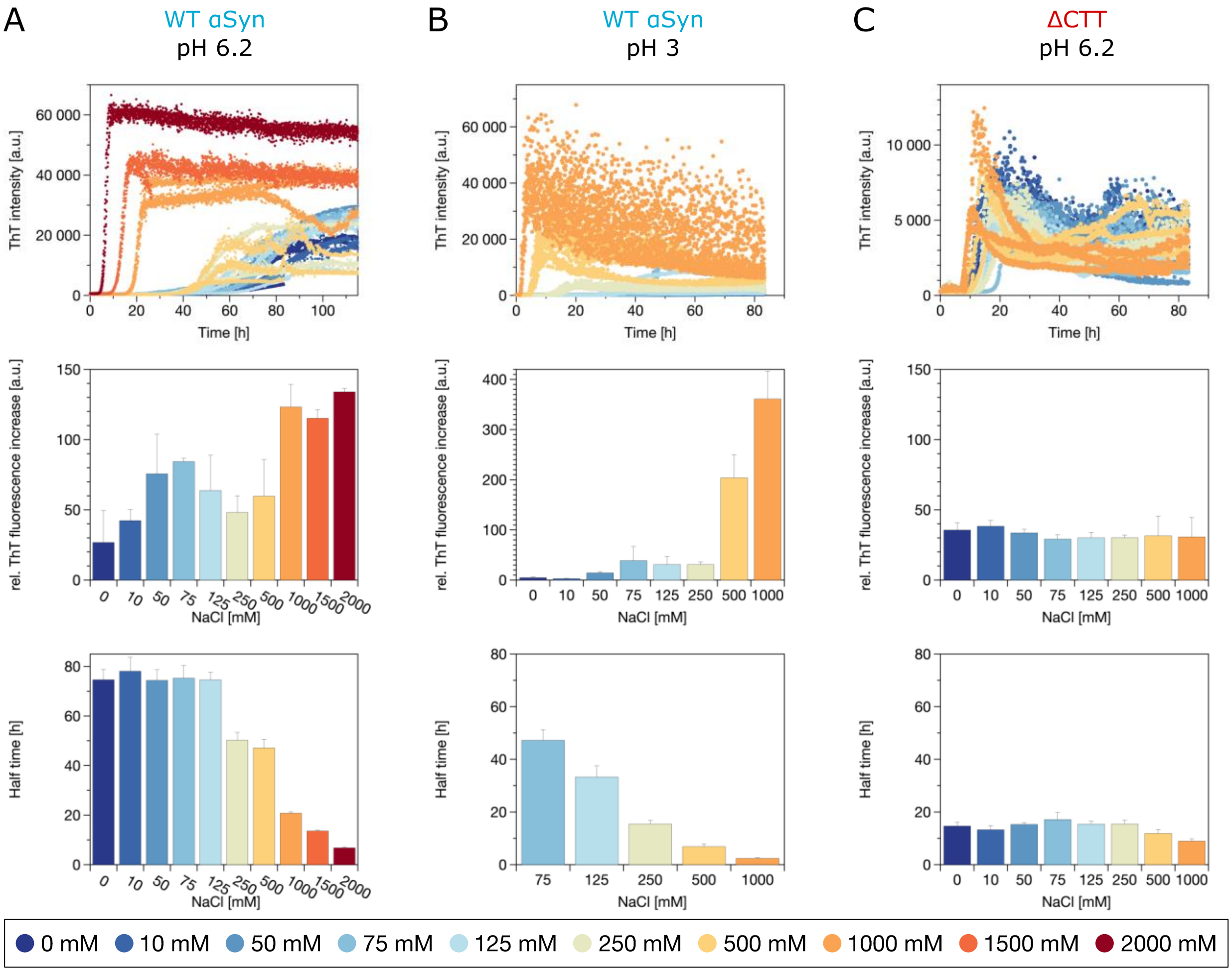


**Fig. S7**: Salt-dependent aggregation of of WT αSyn at pH 6.2 (B), and pH 3 (C) and ΔCTT at pH 6.2 (D) in the absence of crowding agents. From top to bottom panel: ThT-fluorescence aggregation kinetics, rel. ThT fluorescence increase over the time course of aggregation and the half time of aggregation.

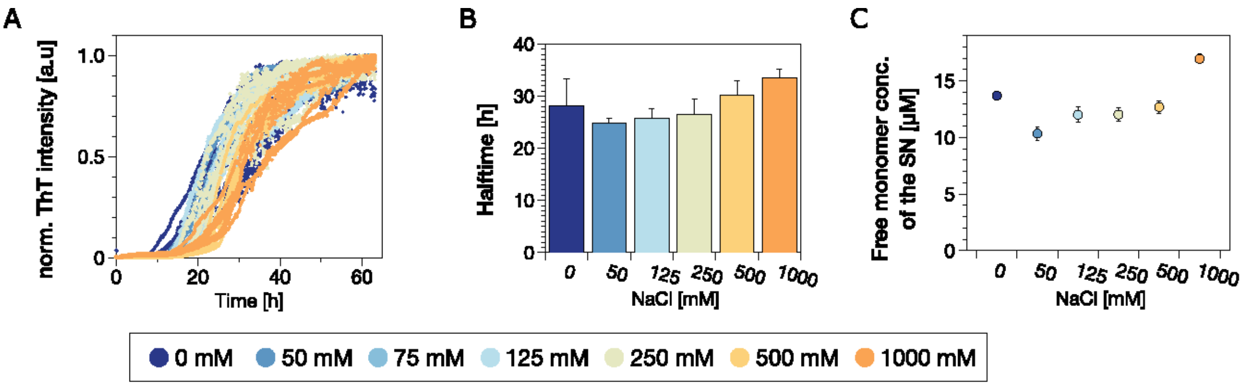


Fig. S8: A) Salt dependent aggregation of 40 µM WT αSyn in the presence of different salt concentrations under crowding (20% PEG400) and quiescent conditions at 25°C (B) with the corresponding half times. C) After 65 hours of incubation, the samples were centrifuged (17 k xg) for 90 min and the concentration of the supernatant was determined.

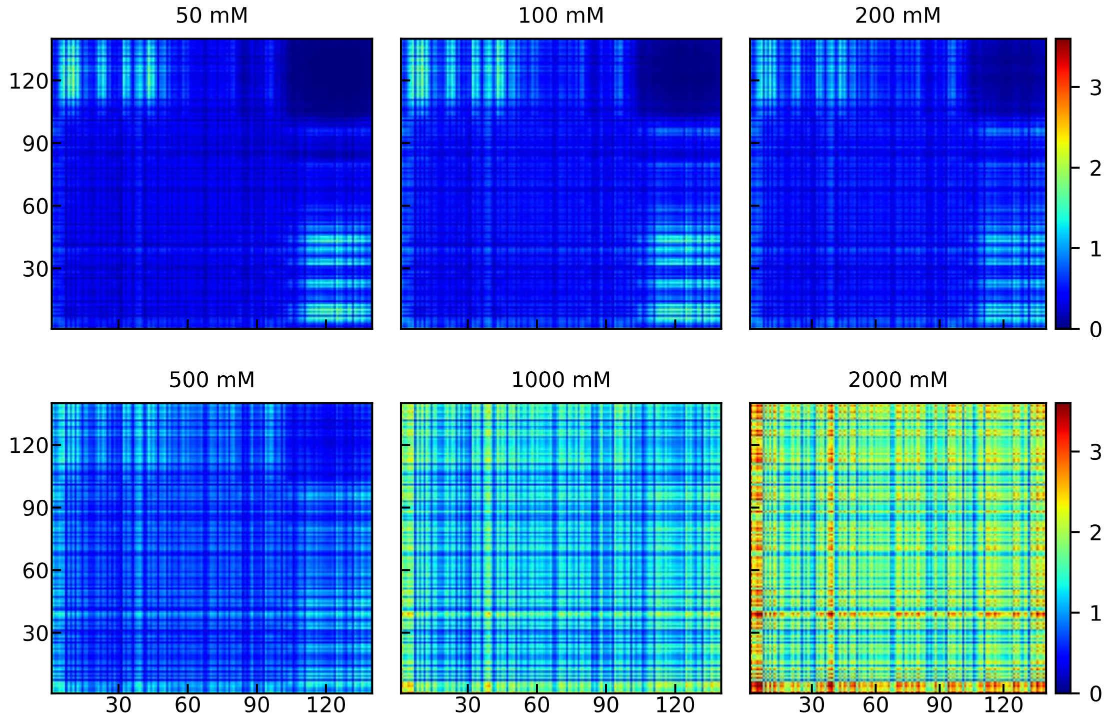


**Fig. S9**: Number of intermolecular contacts (color bar) between pairs of amino acids from the coarse-grained simulations at ε=0.15 kcal/mol (comparable to 20% PEG). The salt concentrations are shown in the title.

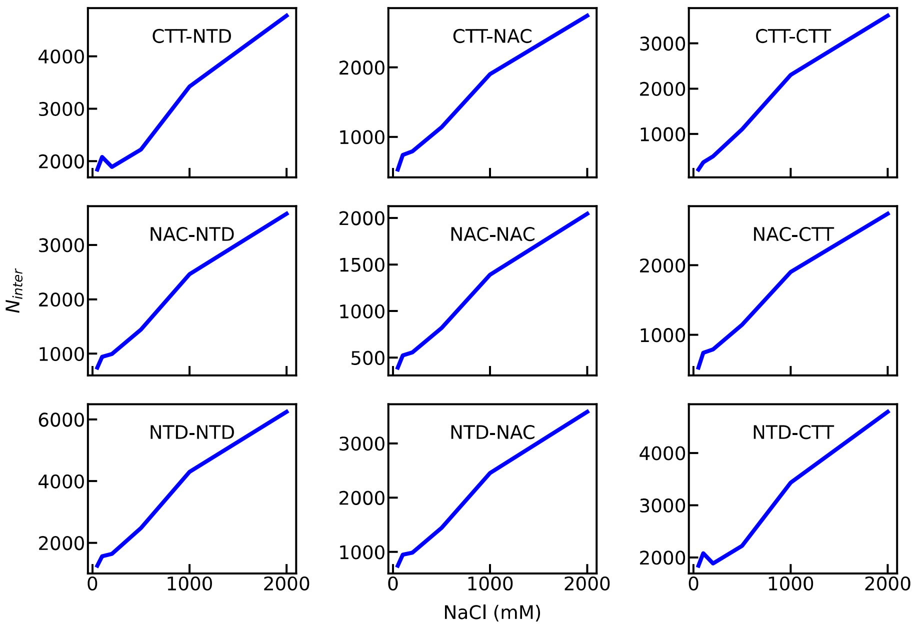


**Fig. S10:** Number of intermolecular contacts between different regions of αSyn from the coarse-grained simulations at ε=0.15 kcal/mol (comparable to 20% PEG). The regions are shown in the legend.

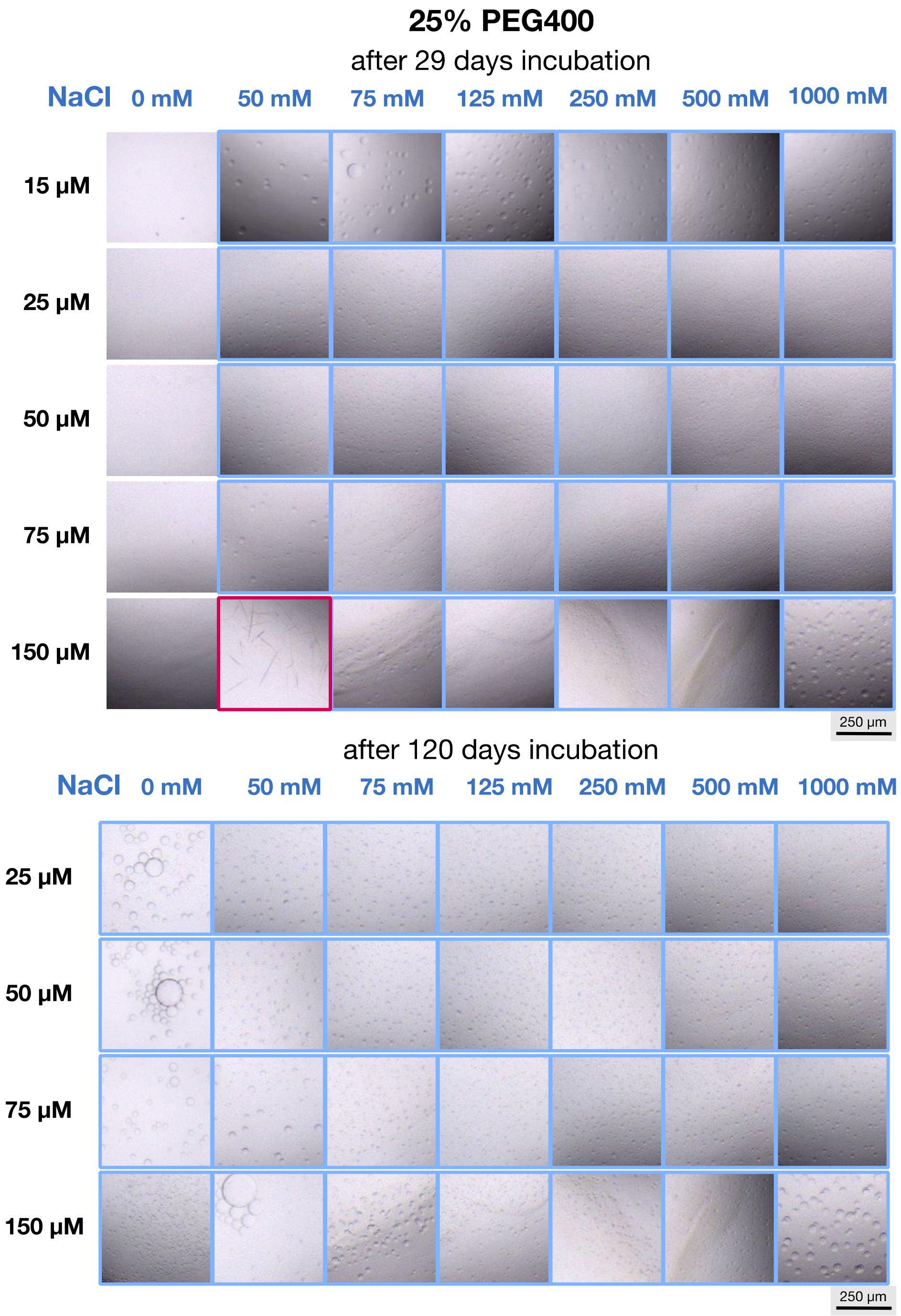


**Fig. S11**: Micrographs (500 x 500 µm) of LLPS formation of WT α-Syn at pH 7 at different salt and protein concentrations after an incubation of 29 days (top) and 120 days (bottom) under crowding conditions (25% PEG400). Droplets are indicated with blue and fibrillar structures with red squares.

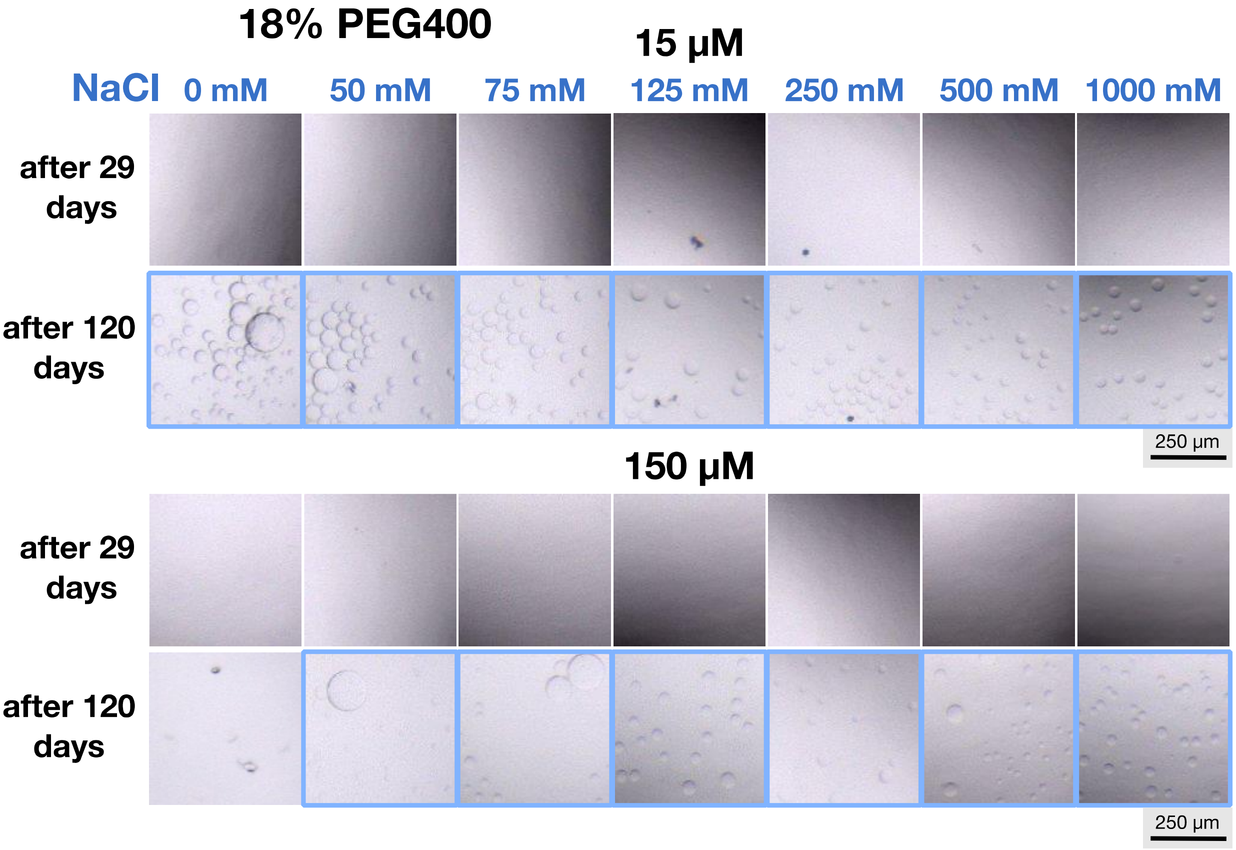


**Fig. S12**: Micrographs (500 x 500 µm) of LLPS formation of 15 µM (top) and 150 µM (bottom) WT αSyn at pH 7 at different salt concentrations under crowding conditions (18% PEG400). Droplets are indicated with blue squares.

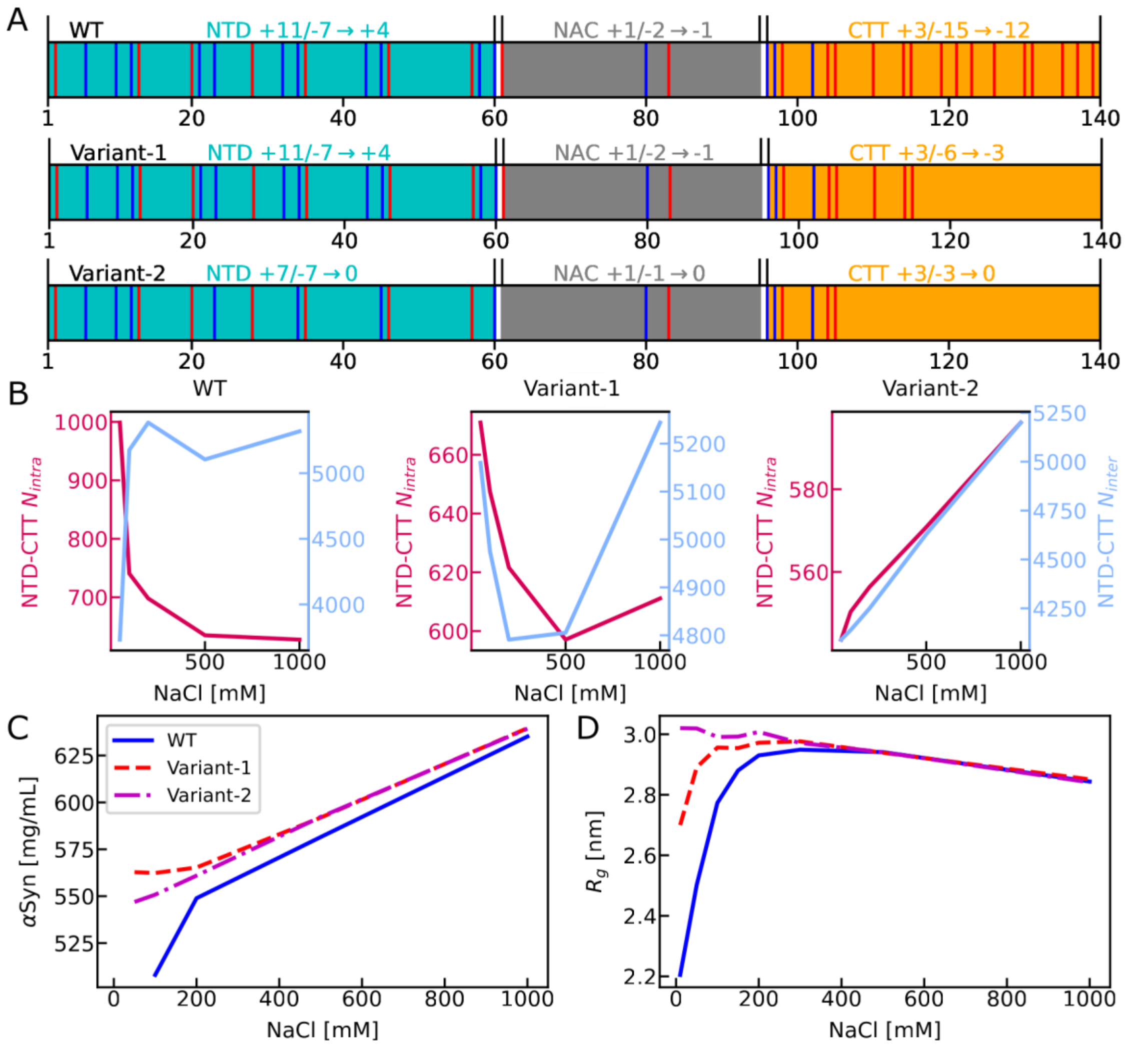


**Fig. S13**: A) Distribution of charged amino acids in wild type (WT) and two variants of αSyn sequence, with positively charged amino acids highlighted in blue and negatively charged ones highlighted in red. The N-terminal domain (NTD, residue 1 to 60) is colored in cyan, non-amyloid-β component (NAC, residue 61 to 95) colored in grey and the C-terminal tail (CTT, residues 96-140) colored in orange. B) The intra- and intermolecular NTD-CTT interactions (ε=0.17 kcal/mol) and C) concentration in the droplet state for the three variants of αSyn from the slab simulation (ε=0.17 kcal/mol). D) $R_g$ from the single-chain simulation (ε=0.15 kcal/mol).

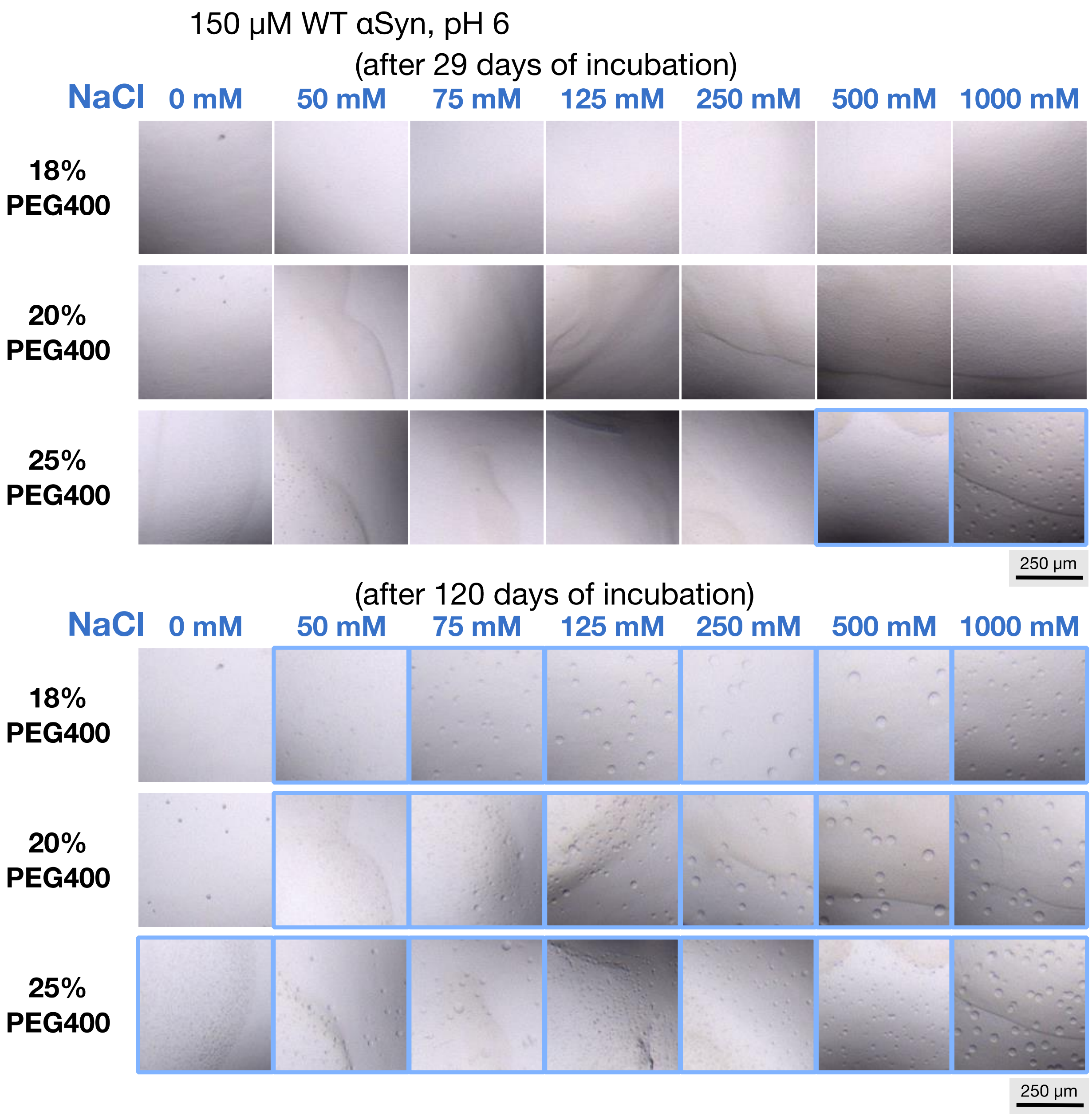


**Fig. S14**: Micrographs (500 x 500 µm) of LLPS formation of 150 µM WT αSyn at pH 6.2 at different salt and PEG400 concentrations after an incubation of 29 days (top) and 120 days (bottom). Droplets are indicated with blue boxes.

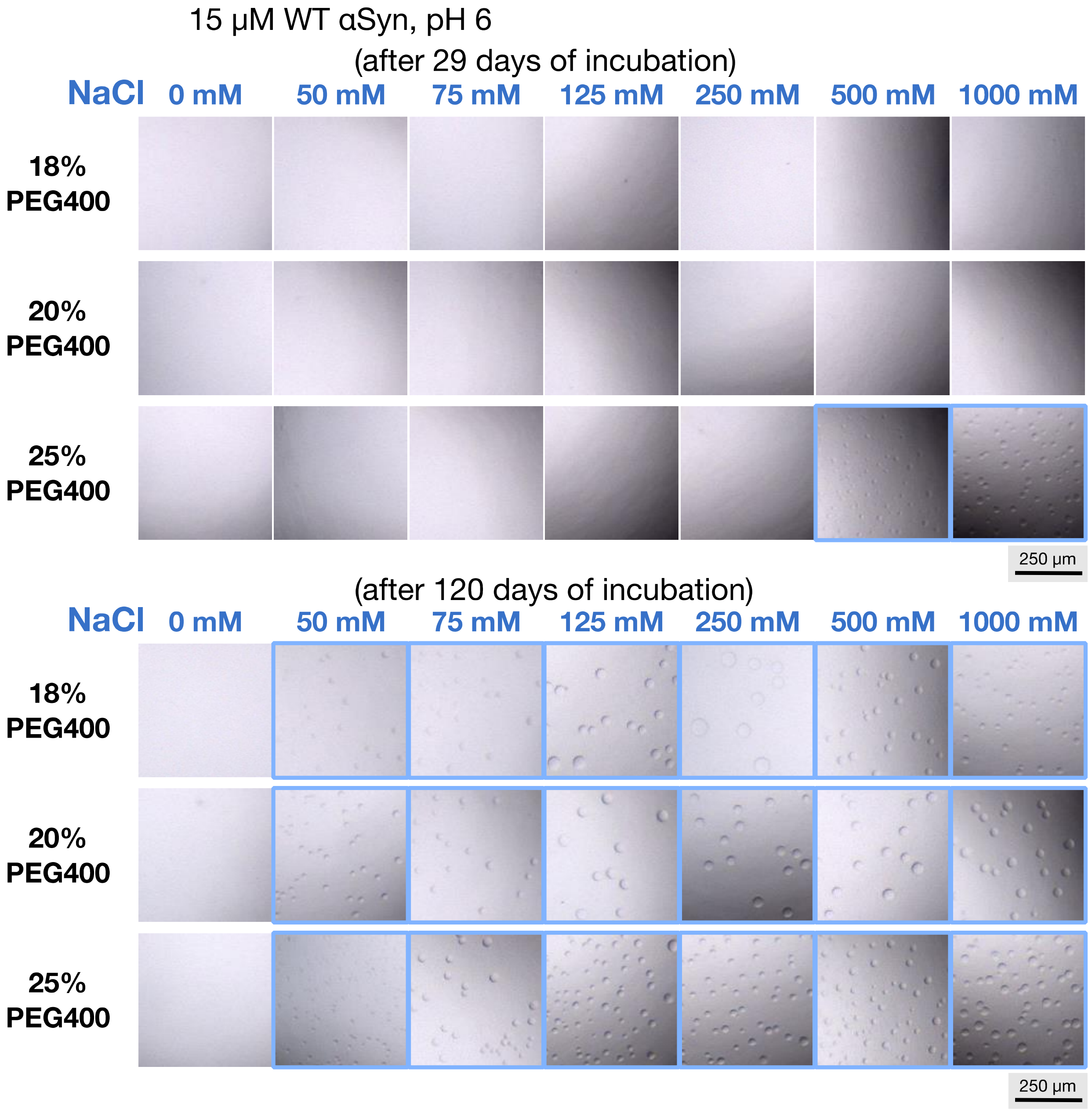


**Fig. S15**: Micrographs (500 x 500 µm) of LLPS formation of 15 µM WT αSyn at pH 6.2 at different salt and PEG400 concentrations after an incubation of 29 days (top) and 120 days (bottom). Droplets are indicated with blue squares.

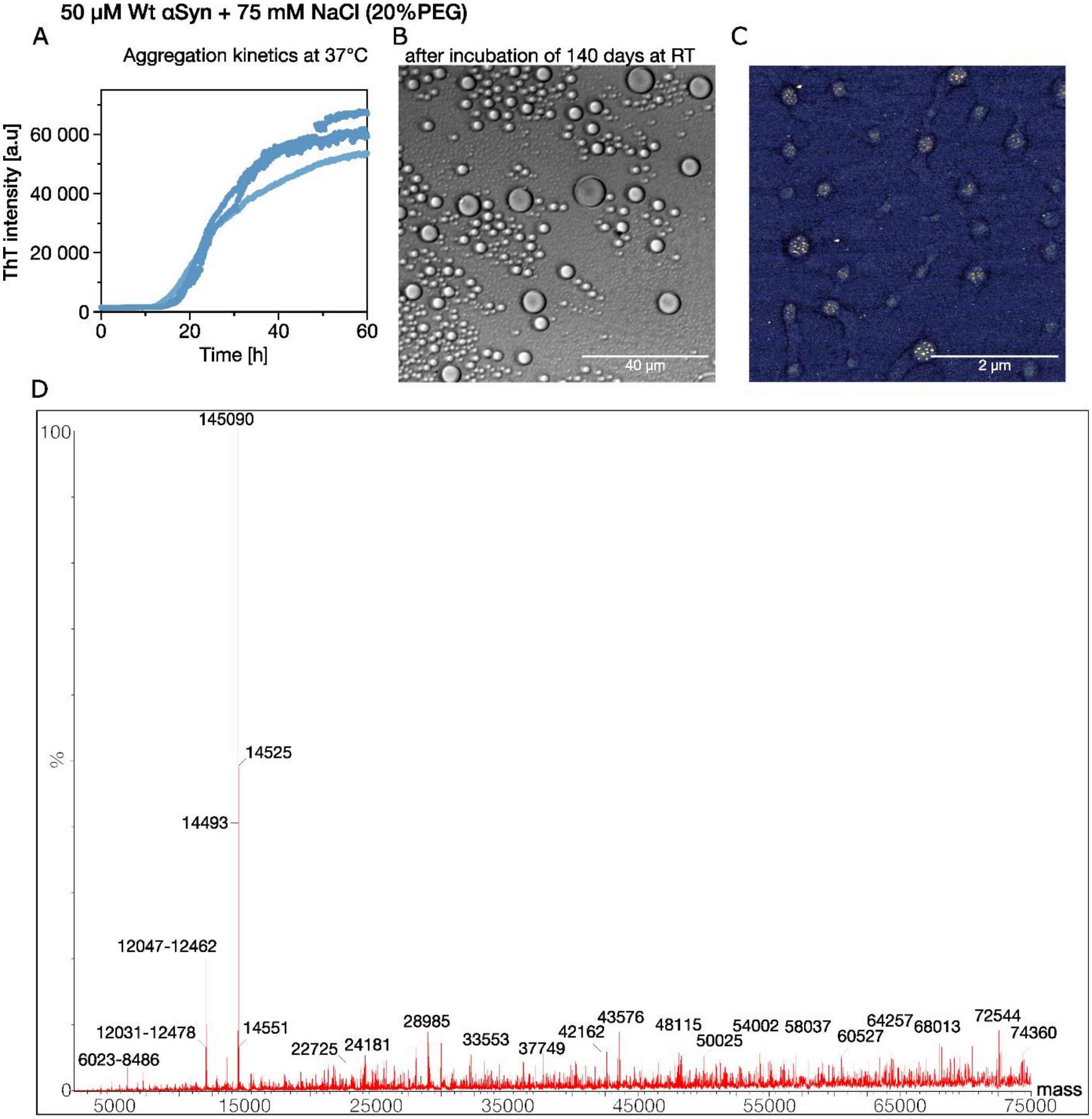


**Fig. S16**: A) 50 µM WT αSyn aggregation in the presence of 75 mM NaCl and 20%PEG400 at pH 6.2 at 37°C was followed by ThT-incorporation experiments. After aggregation the sample was kept in the plate at RT for 140 days. The presence of droplets could be observed by B) TIRF microscopy and C) AFM. D) ESI-TOF LC-MS analysis showed monomeric WT αSyn as the dominant species and no strong degradation over the time course of 140 days.

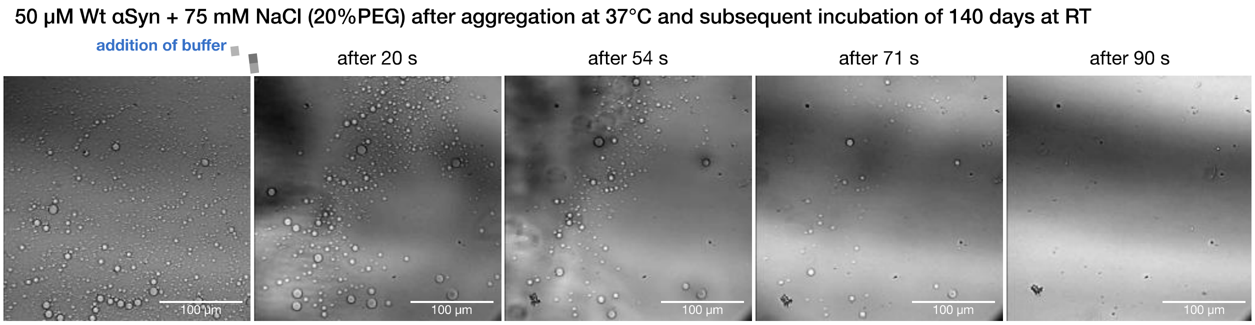


Fig. S17: The sample described in Fig S16 (50 μM WT αSyn in the presence of 75 mM NaCl and 20%PEG400 was kept after aggregation for 140 days at RT) was imaged using TIRF microscopy. The sample was diluted with buffer (1:3) and observed over time. After addition of buffer, most of the droplets disappear within 90 seconds.
